# STRESS-INDUCED GENETIC CHANGE IN FLAX REVEALS GENOME VARIATION MECHANISM

**DOI:** 10.1101/843763

**Authors:** Xiang Li

## Abstract

Inherited genetic change can happen in flax (*Linum usitatissimum*) due to environmental stress. The change can result in different phenotypes in flax progeny. The genetic changes can be induced during one generation and can result in stable genotypes in the next generation. Also the genetic changes are precise and repeatable that homozygous individuals show the same genome reorganization at specific sites in their progeny. Therefore, the genetic re-arrangements are not the result of a random process but a preference for a particular DNA structure, which indicates that possible new genes or functional fragments are formed during this process. The genomes of different varieties of flax are compared to explain the detail of the rapid but intentional genomic changes and regions of variation identified by whole genome comparisons. Possible mechanisms and the potential causes of these rearrangements bring a new light to modern plant genome studies and molecular evolution research.

## Introduction

### 1. Linum usitatissumum

Plants species often show special characteristics in growth, reproduction and nutrition absorption to quickly adapt to their new environment (Saxena et al., 2014). Genomic arrangement can be dynamic in response to stress growth conditions in Flax (*Linum usitatissimum*) (Walbot and Cullis, 1985). Flax is an important plant that has been widely cultivated for commercial use such as for its fiber, linseed oil production and for medicinal purposes (Cullis and Cullis, 2019a). Also, it has many good characteristics that make it a great lab model. First, it has many visible external difference between individuals including the height, weight, seed number, amount of seed, capsule septa hair number and branch number, which makes it easy to distinguish different phenotypes for genome studies. Second, flax is a self-pollinated annual species with a diploid number of 30 chromosomes (Hall et al., 2016. pp.164). This almost obligate inbreeding feature means flax varieties are usually homozygous if there is no stress-caused change or mutation. Thus, differences between generations in characteristics cannot be due to the segregation of pre-existing variation, and narrowing down that the variation is newly formed. Third, frequent genome changes can happen in flax in just one generation in response to nutritional stress (Durrant,1962). The rapid changes in the genome also make flax as a great model to study plant genome variation. Understanding the flax genome variation can help to understand genome reorganization mechanisms, find potential new functional genes or function of sequences, that may possibly be used to improve the agronomic properties of crop species and modify the modern theory and thinking of molecular evolution.

### 2. Stress influence on plant genome

Sessile organisms like plants interact with their environment closely and abiotic and biotic stress can play a key role influencing the plant growth (Takashi, 2010). Biotic stress is caused by the living organism around the plants like bacteria, weeds or insects. Abiotic stresses are other environmental factors like nutritional conditions, temperature, light, drought and salinity. Those stresses seriously influence plant growth (Rejeb, 2000). Multiple studies have found stress-caused plant genome rearrangements. Lucht (2002) states that pathogen can stimulate somatic recombination in *Arabidopsis* and the recombination rate can also be increased when other chemical molecules lead to the defense mechanism in Arabidopsis.

### 3. Flax genotrophs and stress-induced inherited changes

A genotroph has been defined as a stable line derived from *Stormont cirus*, a type of (inducible) fiber flax, following a single generation growth under inducing conditions (Durrant 1962). The *Stormont cirus* variety, also termed a plastic (Pl) variety, has tall apically dominant stem with few branches, which make it good for fiber production. Alternatively, *Bethune* is a flax variety mainly used for oil production because it has been bred to be highly branched and produce more seed. Rapid inheritable phenotype and genome changes have been found in Pl growing under nutrition stress, however, Bethune’s genome is less responsive to nutrition stress. The large (L) genotroph was the progeny of Pl growing in high nutrition, while the small (S) genotroph refers to offspring of Pl growing in low nutrition conditions. L and S are phenotypically and genotypically different from Pl, but remain stable (like *Bethune*) to subsequent generations of growth under inducing conditions (Figure 1). Pl, under non-inducing growth conditions can still remain as an inducible Pl genotroph. Both L and S are true breeding and uninducible to nutrition stresses, and Pl, when grown in a non-inducing environment remains inducible. All the three genotrophs show obvious height differences (Cullis and Cullis, 2019c). However, L6, derived from the large genotroph self-pollinated for 6 generations under low temperatures, is more similar to S compared to L (Durrant, 1971) (The genotroph living in warm greenhouse has kept genome stable for 50 generations). There are many differences in the phenotype between L and S including height (L is taller than S), weight (L is heavier than S), total DNA content (L has more than S), seed capsule septa hair number (L is hairless but S is hairy) (Cullis, 1981) and other genome differences including ribosomal DNA (rDNA) content (L has more rDNA than S), isozyme band pattern (Cullis, 1976), simple sequence repeats (Bickel, 2011) and some characterized DNA polymorphisms. When L was crossed with S, the progeny showed a wide range of height, weight and other phenotypes (Cullis, 1979). Other genotrophs previously described include S3, the third generation from S grown at low temperature and L3 is the third generation from L grown at low temperatures; S6 and L6 are next third generation from S3 and L3, again grown at low temperature. C1, C2, C3 refer to plants derived from Pl but living in good, medium and low nutrition (Cullis 1979, Figure 2). The relationships between the genotrophs is showed in Figure 3.

**Figure 1.**
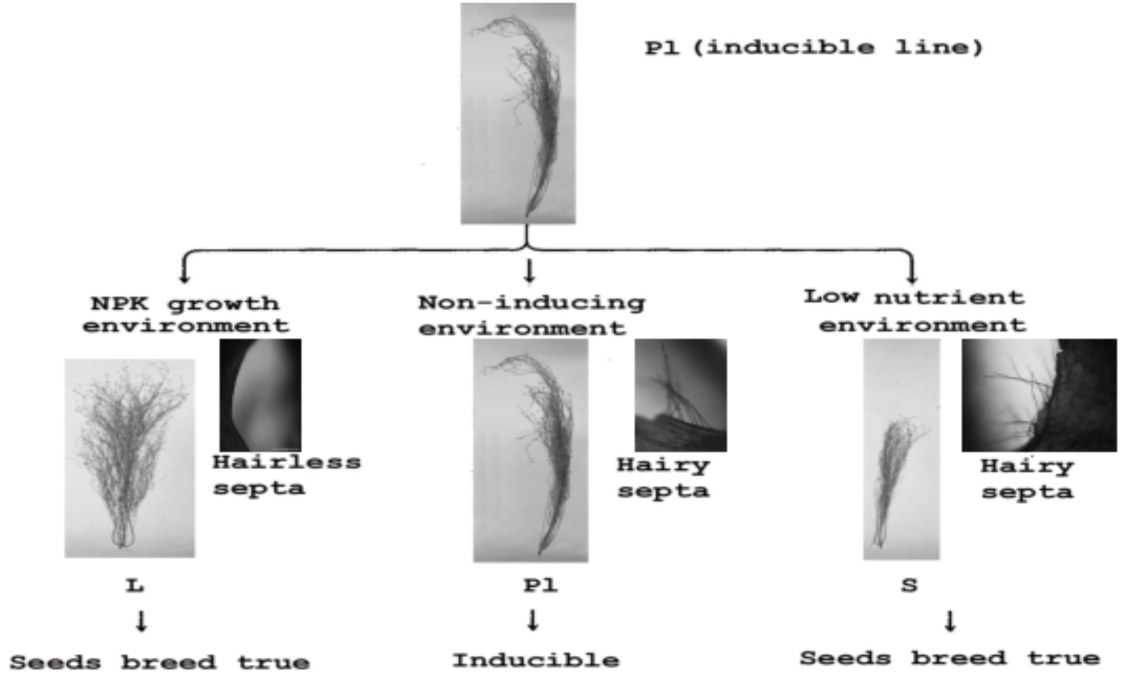
Plant genotroph derived from Pl. Figure provided by C.A. Cullis.

**Figure 2.**
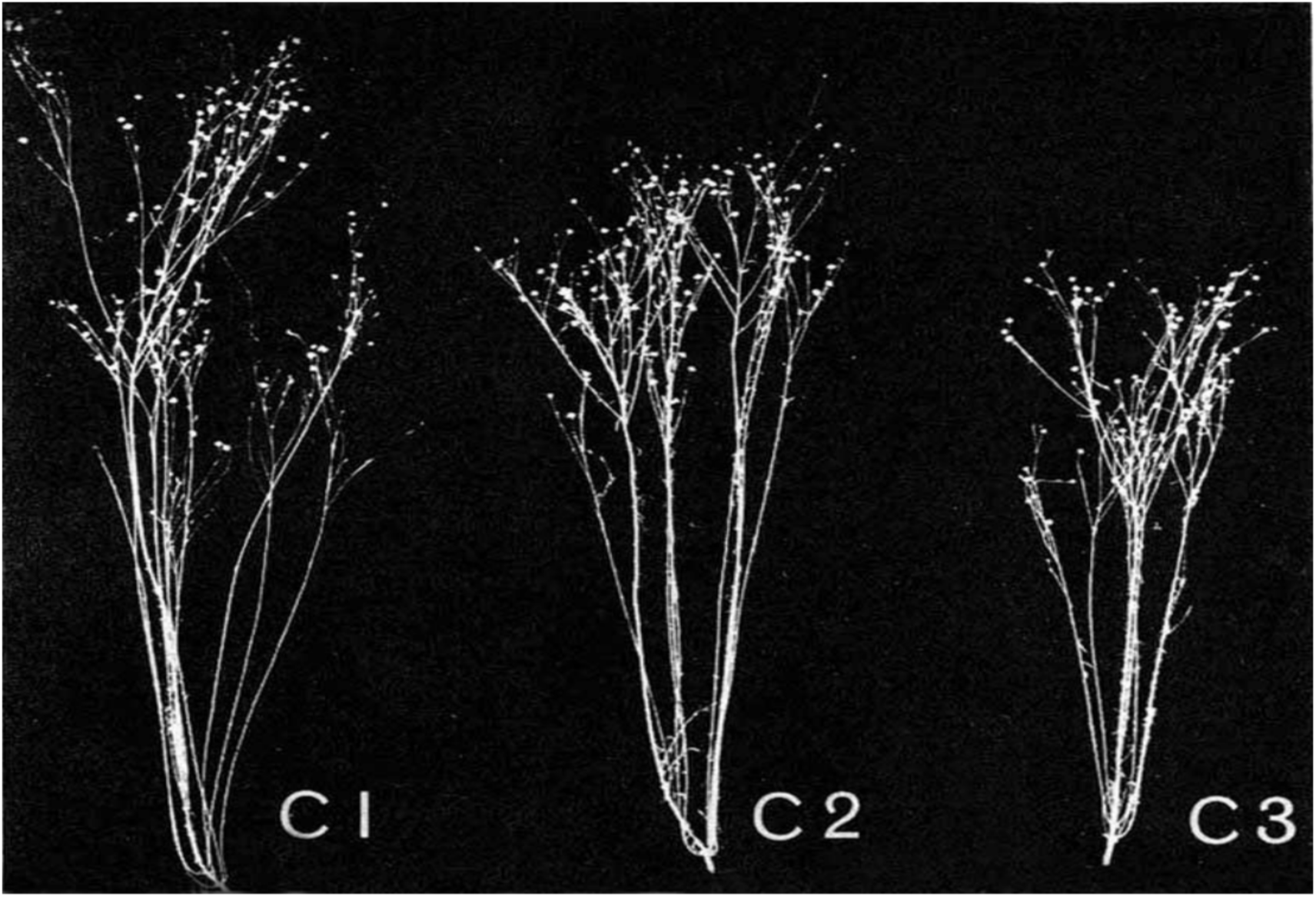
height comparison between C1, C2, C3. Cited from Cullis (2019).

**Figure 3.**
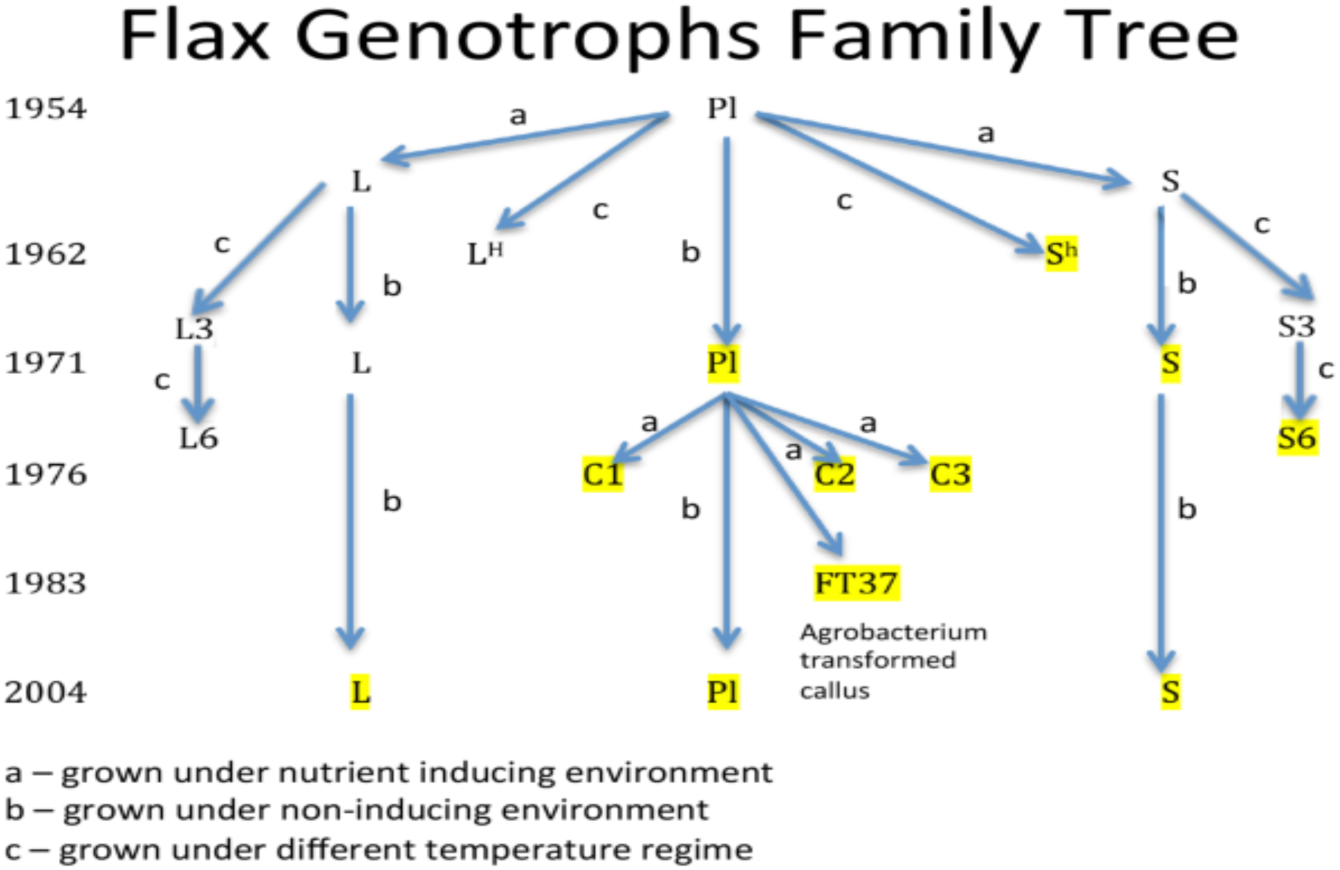
Flax genotroph family, cited from Cullis and Cullis 2019a. The highlighted lines are those for which an Illumina whole genome sequence of at least 20X has been obtained.

### 4. Flax genome polymorphism

The polymorphisms within a plant species can be of many different types such as structural variation (SVs) like insertions, duplications and deletions of large DNA regions (about or more than 1kb), transposable elements (TEs), single nucleotide polymorphisms (SNPs), copy number variations (CNVs) and simple sequence repeats (SSRs) (Saxena, et al., 2014). All of above have been studied and compared between different genotrophs of flax, in an *Agrobacterium tumefaciens* transformed cell line. Some of these polymorphisms have been used as markers to differentiate the genotrophs.

#### SNPs

It has been shown that here are regions containing SNPs when whole genome comparisons are made between Pl and the genotophs (Cullis and Cullis, 2019b). These SNPs don’t show a random distribution but the majority of the genome is maintained constant between Pl and the genotrophs (Cullis and Cullis, 2019b).

#### SVs

The structural variants (SVs), an example of which is the LIS-1 (*Linum Insertion Sequence 1*) insertion caused by nutrition stress, can happen at many sites in the entire genome of plant. LIS-1 is a newly identified 5.7kb insertion in S genotroph that is induced by low nutrition conditions (Chen et al. 2005; Bickle, 2012)(Figure 4). It can be induced in Pl or progeny of Pl✖*Bethune* when grown in low nutrition. The sequence of LIS-1 insertion is not found intact in Pl, B or L genome, except some short matches to both the *Linum* EST database and assembled genome, indicating the LIS-1 is probably a new intentionally assembled fragment inserted into this specific single copy site. Moreover, LIS-1 is not likely the result of a developmental process when comparisons between the control plant individual with the individual living in stress environment are made (Chen et al., 2009). Thus, it is possible that the low nutrition environment directionally triggers this insertion to be made. Different sets of primer pairs have been designed to detect the LIS-1 insertion based on the figure 1. It showed that when LIS-1 is completely inserted, the 2/3’ and 18a/19’ will amplify a fragment, however, 2/pl9 will lose the band due to inadequate elongation time for 5kb band (Cullis, 2017). Although, the sequencing of LIS-1 is done and the apparent partial transposon signature at the insertion site (Bickel, 2012), how this insertion happens, how flax controls this arrangement, and what are the functions of this insertion, are not well known.

**Figure 4.**
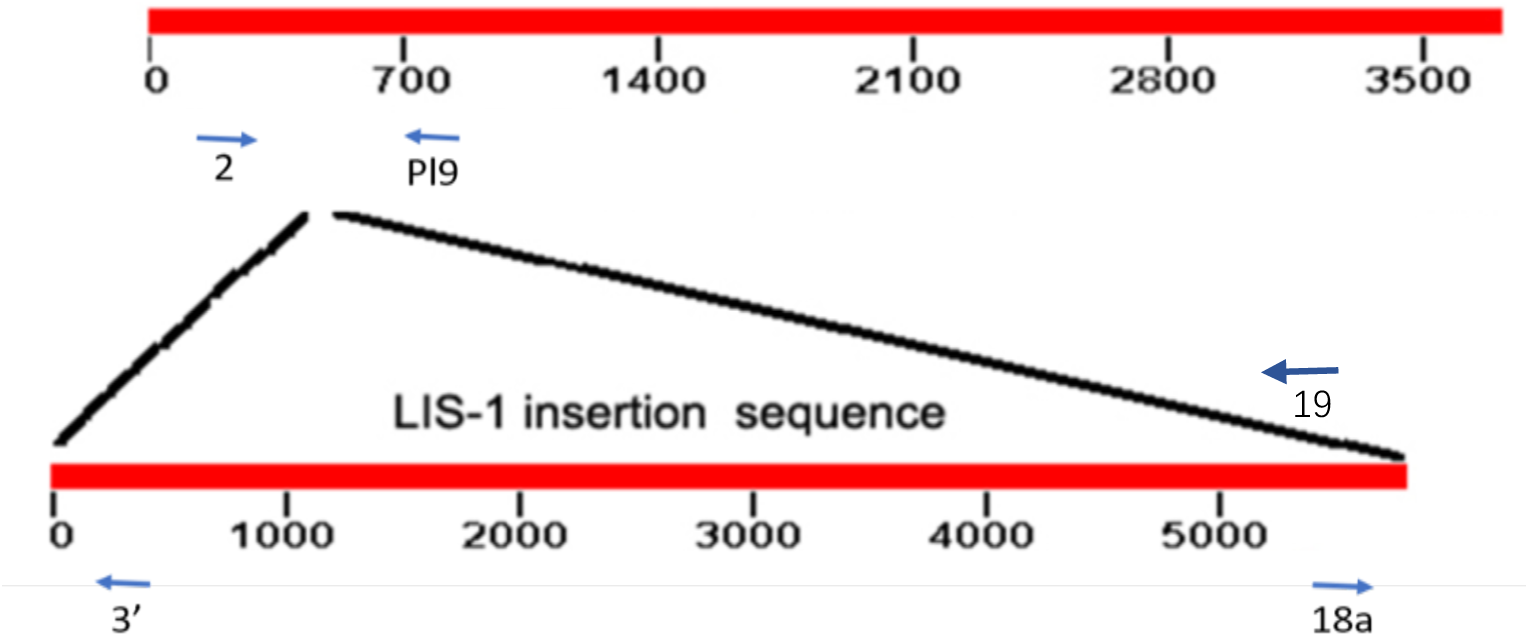
LIS-1 insertion model.

#### rRNA and total amount of DNA

Ribosomal RNA (rRNA) is essential for forming the ribosomes and for protein production. It has been found that L line had more rRNA coding sequences (rDNA) than S line and the percentage of rDNA in the genome of L is nearly 50% more than rDNA percentage in the S (Cullis, 1975, 1976). There are also large total DNA content differences between different genotrophs. Cullis (2019b) stated that *Bethune* and Pl have different genome sizes, with the genome of Pl nearly twice of *Bethune*. Otherwise, around 30% of Pl genome sequence are not present in the *Bethune* reference sequence (Cullis and Cullis, 2019b). When the L genotroph is compared with S, L had 16% more nuclear DNA than S with the Pl genotroph having an intermediate amount (Evans, et al., 1966). Another comparison between Pl and C3 shows a reduction of C3 total genome by 6.8% relative to Pl, mainly due to the difference between highly repetitive sequence numbers (Cullis, 2005). This evidence demonstrates a large-scale genome variation between different varieties or different induced genotrophs.

### 5. Plant transposable elements

Transposons were first described in *Zea mays* more than 80 years ago (McClitock et al., 1931), and today they are ubiquitous and known as “jumping genes”. They make up over 50% of nuclear DNA in plants with large and complex genomes and play a key role in evolution and genome variation (Bennetzen, 2000). Transposable elements (TEs) can create DNA re-arrangements for plant rapid adaption. They not only work as mobile fragments to alter the genome function, but also provide the chance to create a new better variety for evolution. There are some markers that identify a transposon element, for example, the DNA transposable elements always come with TIRs (terminal inverted repeats) ranging from tens of bases to hundreds of bases. Also, the DNA sequences at the ends of transposons are usually the same due to the insertion mechanism (Figure 5). When a transposon leaves the duplicated region remains. This duplication has been termed as the transposon footprint. When transposon is excised, the same sequence would get together and form a repetitive sequence like ATGCATGC.

**Figure 5.**
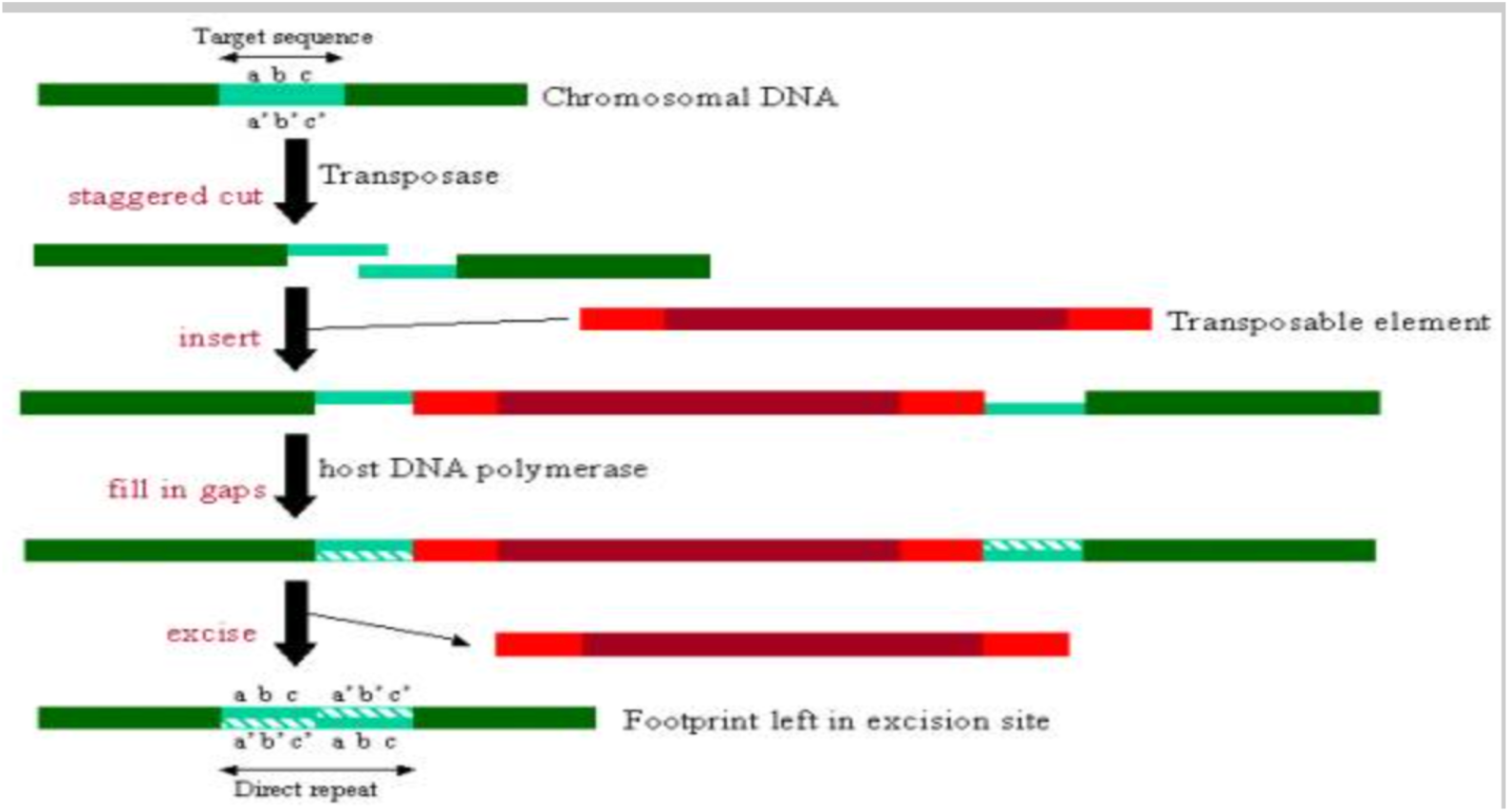
Transposon footprint. When transposable element is excised, the end joints and makes a short repeat sequence. Cited from https://www.ebi.ac.uk/interpro/potm/2006_12/Page1.htm

## Materials and Methods

### 1: Plant material

Three sets of plants have been analyzed.

1. The first set (60 samples) were collected from Pl growing in low nutrient conditions. The leaves were collected at bottom, middle and top of the main stem so each individual provided 3 leaf samples. These samples are labelled as 1T, 1M, 1B, the number indicates the plant and T, M, B refer to top, middle and bottom of the main stem. The sample 12 has two 12 top samples, one of which was collected from the growing tip.
2. The second set of plants are inducible Pl living in the low nutrient environment. Leaves were collected from different branches. The samples are labelled as 1.1, 1.2, 2.1, 2.2, where the first number indicates the specific plant and the second refers to branch. There can be more than 5 branches in one individual and all branches were photographed.
3. The third set are DNAs from Pl treated either with npk (n: 1 percent ammonium sulphate; p: 1 percent solution of triple superphosphate; k:1 percent solution of potassium chloride. Durant (1962)) or with water. The samples are labelled as “1nl” for sample 1, npk treated, DNA collected from leaves, or “2wm”, means sample 2, water treated, DNA collected from meristem. All the isolated DNAs were stored at 4 °C. The leaf samples were stored at - 20°C before DNA extraction. The significance of the different environments set is that they can cause visually phenotypical changes in subsequent generations of flax (Durant, 1962), thus, they are useful for flax genome comparison to identify any genetic changes.

#### DNA extraction and quantity test

DNA was extracted from 0.1 gram green leaves by using DNeasy plant mini kit (QIAGEN). The DNA concentration was determined using the Qbit or nanodrop. The original concentrations are given in table 3.

### 2: IGV (Integrative Genomics Viewer)

The Integrative Genomics Viewer (IGV) is a high-performance visualization tool for interactive exploration of large, integrated genomic datasets (version 2.4.4, downloaded from https://software.broadinstitute.org/software/igv/). We used IGV to compare the genomic difference between genotrophs. The IGV alignments mapped many gaps when comparing Pl or C3 with *Bethune* (*Bethune* is used as a reference sequence) that represent large genome variation from either deletions or insertions.

### 3: Polymerase chain reaction (PCR)

The program used for the amplification is in table 1. Primer sequences used are listed below in table 2 and designed by Professor Cullis. The primers were chosen from regions that differed between the genotrophs, either across the insertion region of LIS-1, or where there were gaps in the alignments of whole genome sequencing, identified using IGV, when individuals were compared to reference flax genome. Each reaction contains 25ul solution including 1ul (2-5ng) sample DNA, 1ul primer mix (10mol/L), 12.5ul of Taq mix and 10.5ul distilled water or 25ul Tag, 22ul water, 2ul primer and 1ul DNA for 50ul solution (exact DNA amount is based on the original concentration). After PCR reaction, the samples are stored at 4°C to prevent DNA degradation. QIAquick PCR purification kit was used to clean up amplified DNA prior to sequencing. The concentration was tested using a Qbit.

**Table 1.** PCR reaction protocol. The elongation time can vary when particular fragment size is known to be amplified. When PCR is finished, the DNA will be store at 4 °C.

| Initiation | Cycling: 39 cycles | Final elongation | Store |
| --- | --- | --- | --- |
| 95°C 2minutes | 95°C 30s; 60°C 30s; 72°C 1min | 72°C 2mins | 12°C |

**Table 2.**
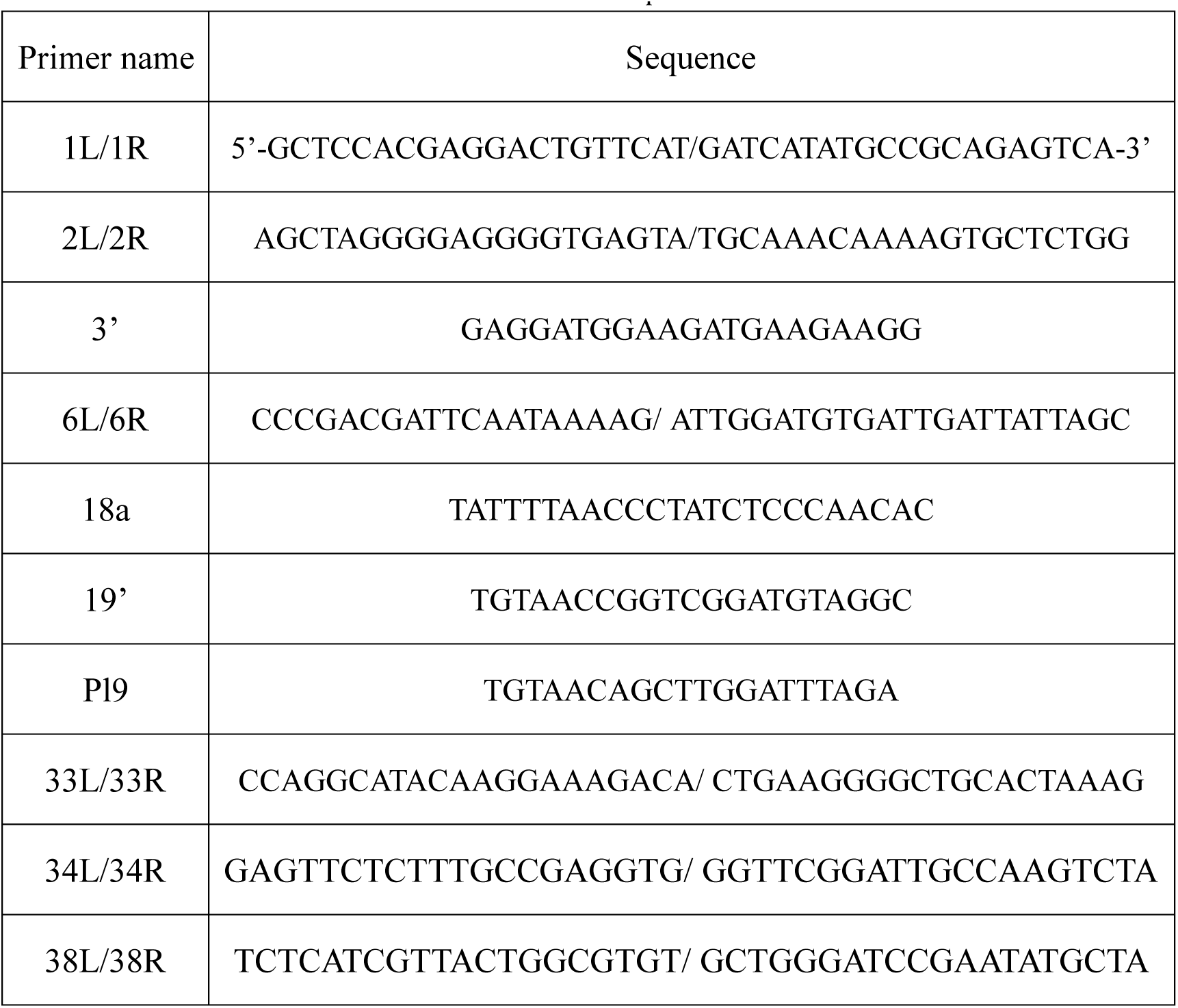
Primer number and sequences used for PCR

**Table 3.**
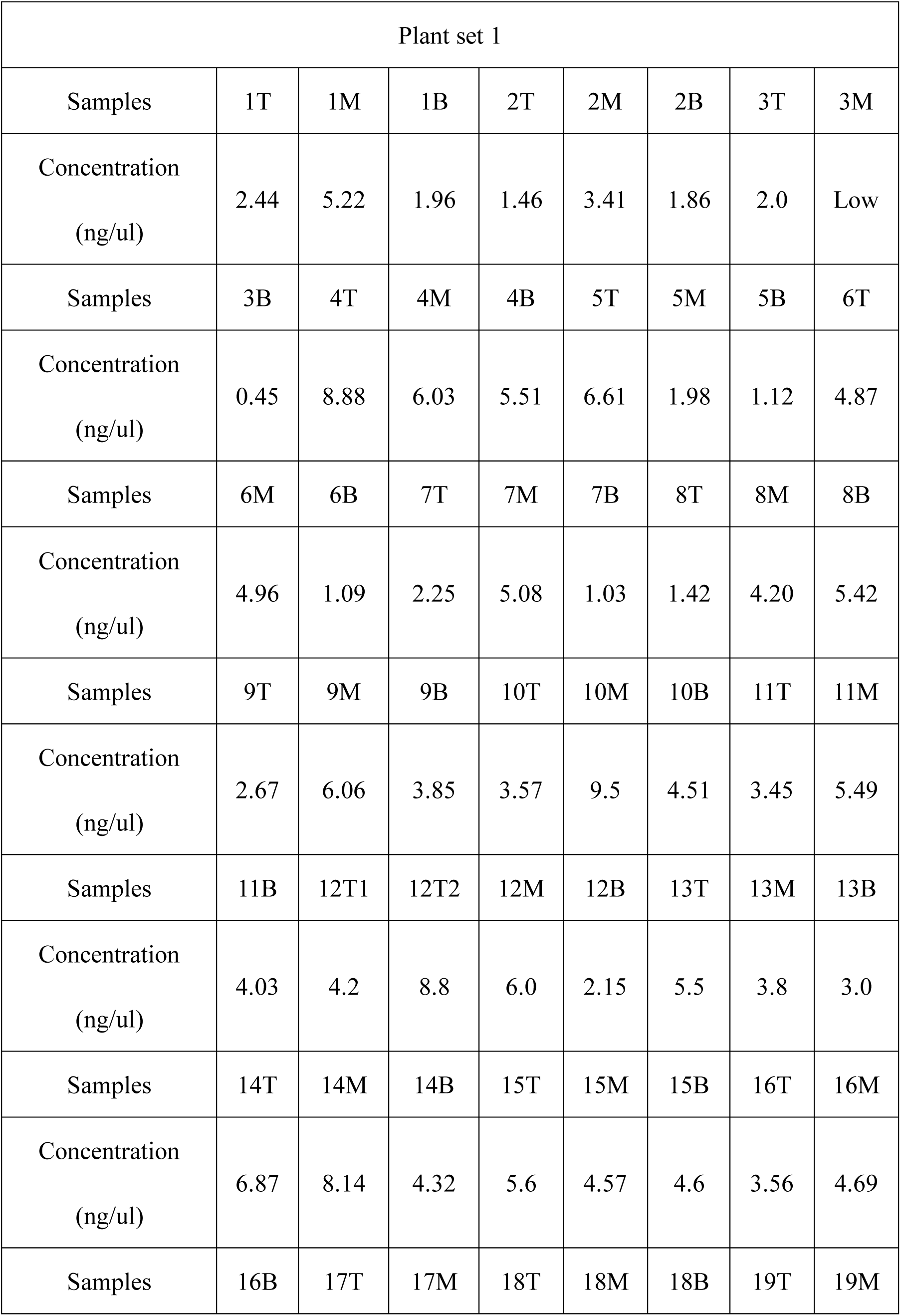
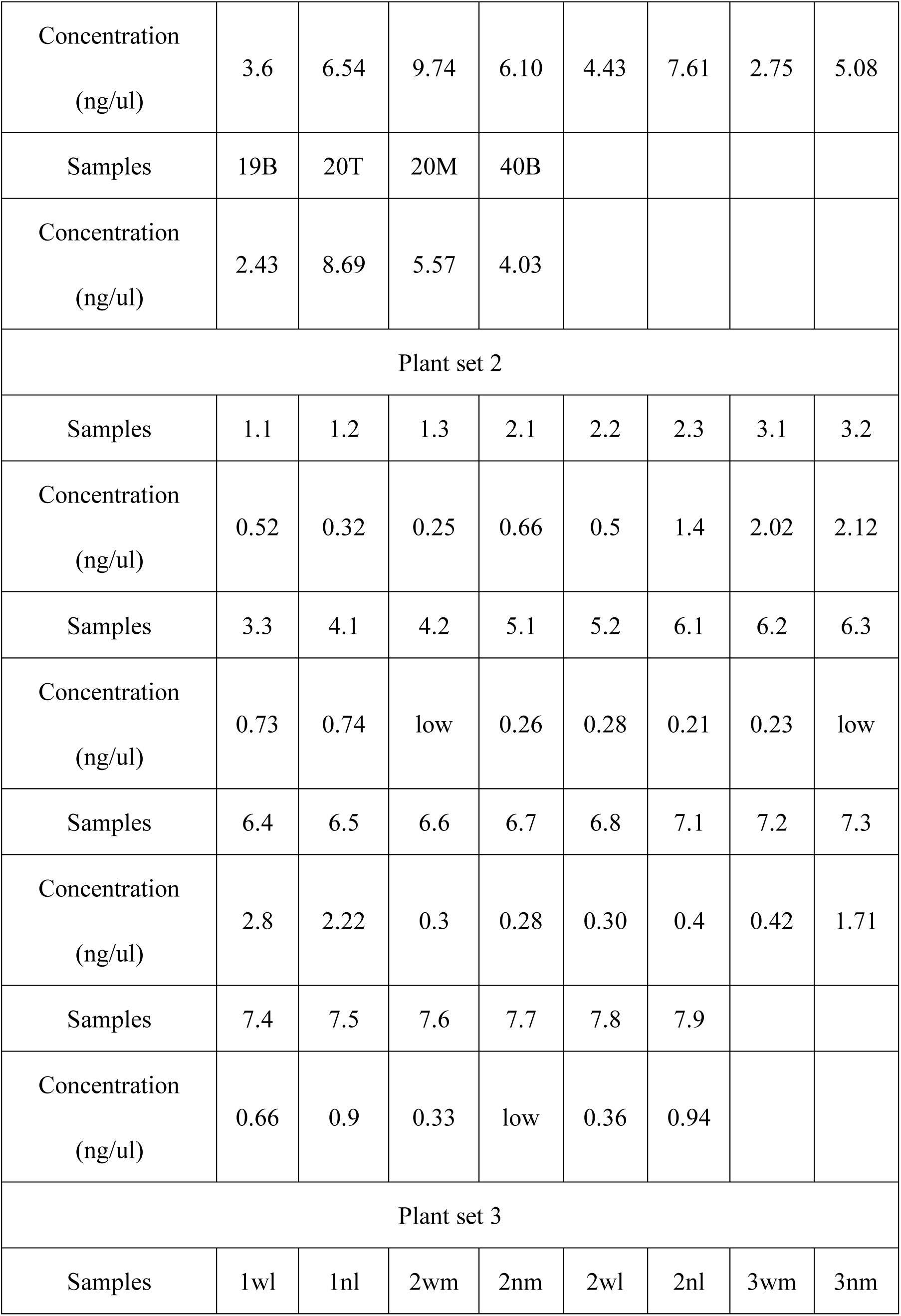
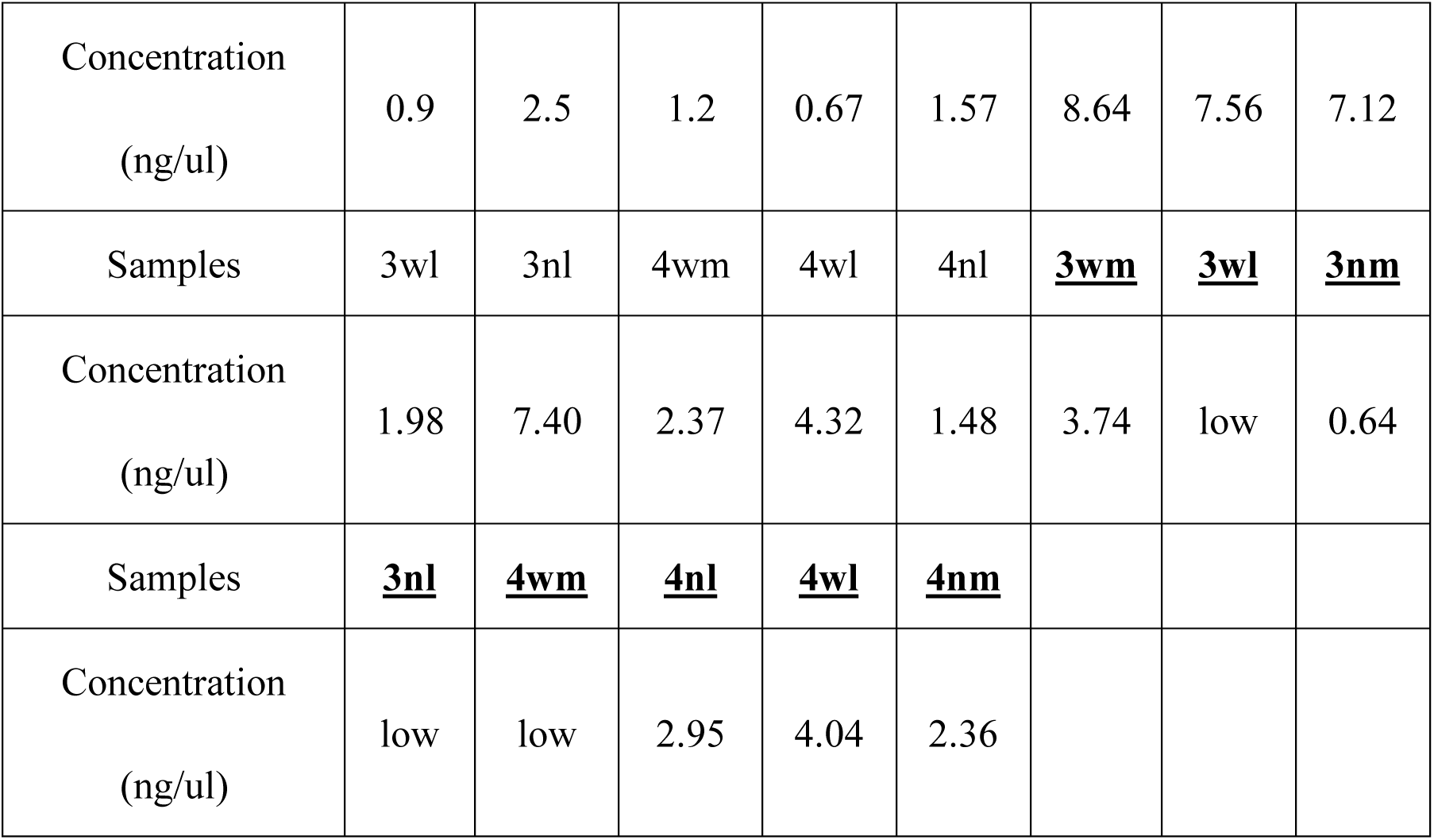
Original DNA concentration

| Plant set 1 |  |  |  |  |  |  |  |  |
| --- | --- | --- | --- | --- | --- | --- | --- | --- |
| Samples | 1T | 1M | 1B | 2T | 2M | 2B | 3T | 3M |
| Concentration<br>(ng/ul) | 2.44 | 5.22 | 1.96 | 1.46 | 3.41 | 1.86 | 2.0 | Low |
| Samples | 3B | 4T | 4M | 4B | 5T | 5M | 5B | 6T |
| Concentration<br>(ng/ul) | 0.45 | 8.88 | 6.03 | 5.51 | 6.61 | 1.98 | 1.12 | 4.87 |
| Samples | 6M | 6B | 7T | 7M | 7B | 8T | 8M | 8B |
| Concentration<br>(ng/ul) | 4.96 | 1.09 | 2.25 | 5.08 | 1.03 | 1.42 | 4.20 | 5.42 |
| Samples | 9T | 9M | 9B | 10T | 10M | 10B | 11T | 11M |
| Concentration<br>(ng/ul) | 2.67 | 6.06 | 3.85 | 3.57 | 9.5 | 4.51 | 3.45 | 5.49 |
| Samples | 11B | 12T1 | 12T2 | 12M | 12B | 13T | 13M | 13B |
| Concentration<br>(ng/ul) | 4.03 | 4.2 | 8.8 | 6.0 | 2.15 | 5.5 | 3.8 | 3.0 |
| Samples | 14T | 14M | 14B | 15T | 15M | 15B | 16T | 16M |
| Concentration<br>(ng/ul) | 6.87 | 8.14 | 4.32 | 5.6 | 4.57 | 4.6 | 3.56 | 4.69 |
| Samples | 16B | 17T | 17M | 18T | 18M | 18B | 19T | 19M |
| Concentration<br>(ng/ul) | 3.6 | 6.54 | 9.74 | 6.10 | 4.43 | 7.61 | 2.75 | 5.08 |
| Samples | 19B | 20T | 20M | 40B |  |  |  |  |
| Concentration<br>(ng/ul) | 2.43 | 8.69 | 5.57 | 4.03 |  |  |  |  |
| Plant set 2 |  |  |  |  |  |  |  |  |
| Samples | 1.1 | 1.2 | 1.3 | 2.1 | 2.2 | 2.3 | 3.1 | 3.2 |
| Concentration<br>(ng/ul) | 0.52 | 0.32 | 0.25 | 0.66 | 0.5 | 1.4 | 2.02 | 2.12 |
| Samples | 3.3 | 4.1 | 4.2 | 5.1 | 5.2 | 6.1 | 6.2 | 6.3 |
| Concentration<br>(ng/ul) | 0.73 | 0.74 | low | 0.26 | 0.28 | 0.21 | 0.23 | low |
| Samples | 6.4 | 6.5 | 6.6 | 6.7 | 6.8 | 7.1 | 7.2 | 7.3 |
| Concentration<br>(ng/ul) | 2.8 | 2.22 | 0.3 | 0.28 | 0.30 | 0.4 | 0.42 | 1.71 |
| Samples | 7.4 | 7.5 | 7.6 | 7.7 | 7.8 | 7.9 |  |  |
| Concentration<br>(ng/ul) | 0.66 | 0.9 | 0.33 | low | 0.36 | 0.94 |  |  |
| Plant set 3 |  |  |  |  |  |  |  |  |
| Samples | 1wl | 1nl | 2wm | 2nm | 2wl | 2nl | 3wm | 3nm |
| Concentration<br>(ng/ul) | 0.9 | 2.5 | 1.2 | 0.67 | 1.57 | 8.64 | 7.56 | 7.12 |
| Samples | 3wl | 3nl | 4wm | 4wl | 4nl | **<u>3wm</u>** | **<u>3wl</u>** | **<u>3nm</u>** |
| Concentration<br>(ng/ul) | 1.98 | 7.40 | 2.37 | 4.32 | 1.48 | 3.74 | low | 0.64 |
| Samples | **<u>3nl</u>** | **<u>4wm</u>** | **<u>4nl</u>** | **<u>4wl</u>** | **<u>4nm</u>** |  |  |  |
| Concentration<br>(ng/ul) | low | low | 2.95 | 4.04 | 2.36 |  |  |  |

### 4: Sequencing

Sequencing was done by Eurofins company with Sanger sequencing method. The sample preparation follow the sequencing guidelines. The ABI trace was viewed with software “4 Peaks” (downloaded from https://nucleobytes.com/4peaks/index.html).

### 5: DNA sequence comparison

Basic Local Alignment Search Tool (BLAST) was used to compare different DNA sequences. Blastn is used for nucleotide blast. The DNA sequence input in BLAST was confirmed by using the sequencing results from both the left and right primers, corrected and completed based on the sequencing trace. Both “somewhat similar” and “high similarity” options were used. The blast result is compared with whole genome sequencing data to see if the SNPs were true and IGV to look at if the result is consistent with the gap size that was identified with IGV.

## Results

The result show the genome comparison of the three different plant sets described in the Materials and Methods. Specifically, samples that are leaves from different height on one branch, leaves from different branches, and leaves from plants grown in different nutritional environments.

### 1. Samples collected from top, middle and bottom of the same individual

The figure 6 shows the amplification of one of the junctions of the LIS-1 insertion by the primer pair 2 and 3’. Two bands that are around 90bp and 200bp are contained in almost every samples, which refers to unknown normal genomic DNA sequences. The expected 500bp fragment covering part of the LIS-1 insertion does not show in every sample, indicating genome variation. Sample 16B has the 500bp band but loses the other two bands, while 9B, 14T, 15T, 15M,19T have all three bands.

**Figure 6.**
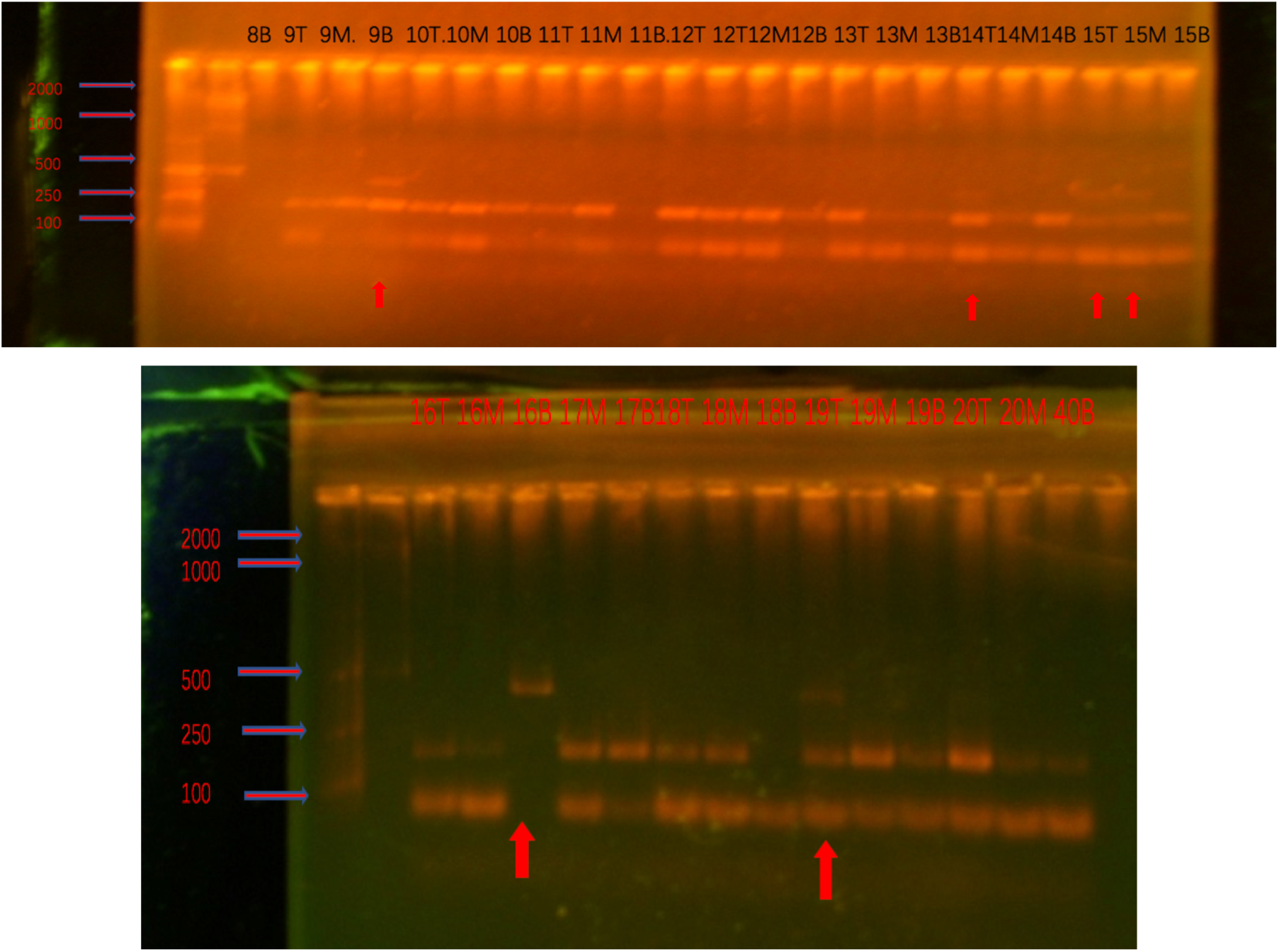
Gel result of PCR reaction with primer 2/3’. 1.2% agarose gel, 120V. The red arrow points out the larger band. T,B and M refer to top, middle and bottom. Bio easy marker I and II are used in the left two lanes. The picture only shows some of the samples because the rest of the bands are too faint to make a conclusion. The summary is in the table 1.

Figure 7 shows the PCR reaction of primer 2/pl9, which covers the left and right end of LIS-1 insertion. All samples show one band around 700-800bp except 4B.

**Figure 7.**
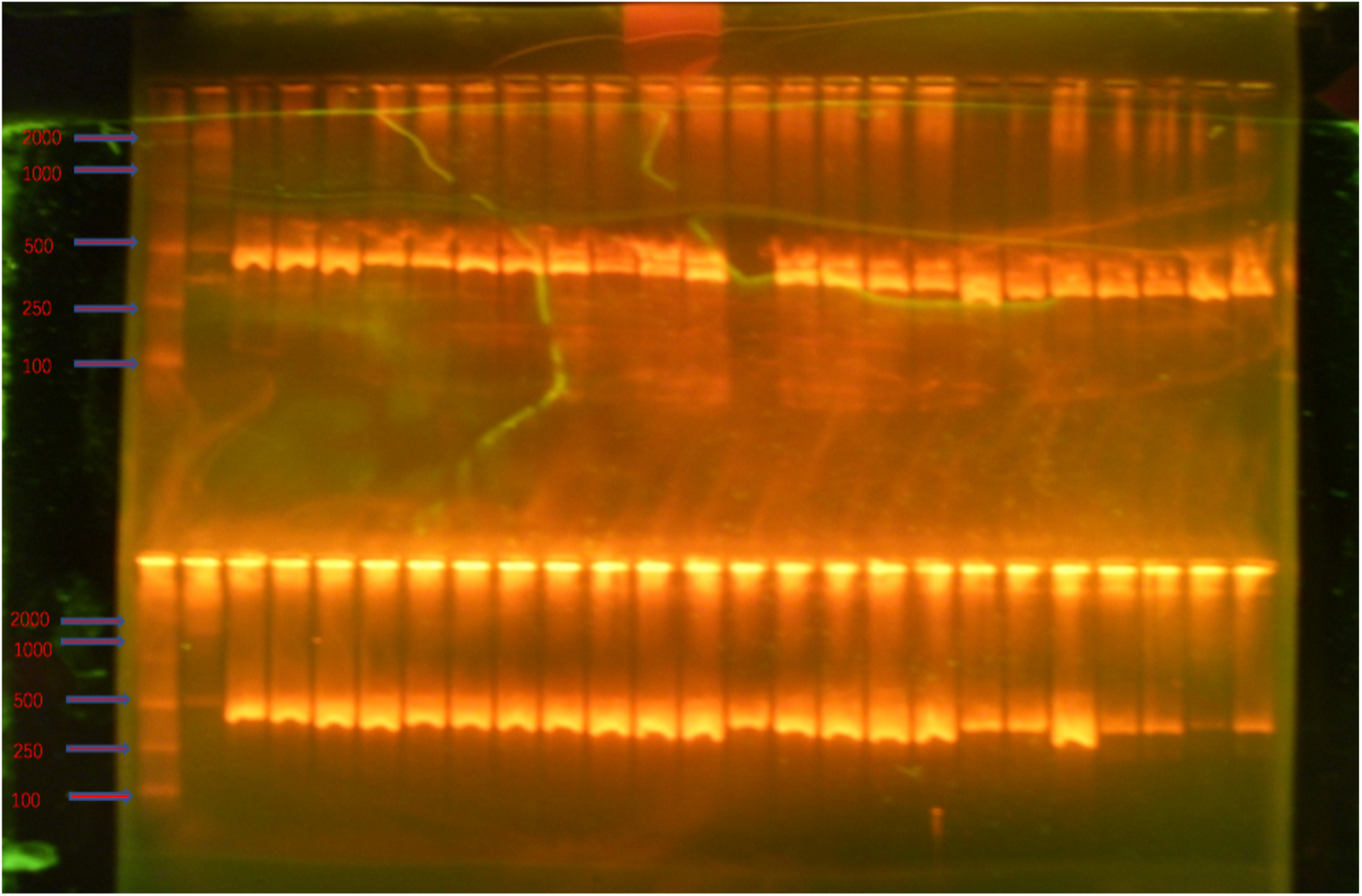
Gel picture of PCR reaction with 2/pl9. 1.2% agarose gel, 1.2V for 2 hours. Bio easy marker I and II are used in the left two lanes.

From the figure 8, 18a/19’ covers another end of LIS-1 insertion. The extra band indicates the insertion formed in terms of right end of LIS-1. The sample 3B and 6B shows the extra band while the 3M,3T or 6M,6T didn’t show the band, which can suggest that variation can change in different stages. But, the 7T and 2M show that the extra band can happen in any leaves and don’t show a bottom-up influence.

**Figure 8.**
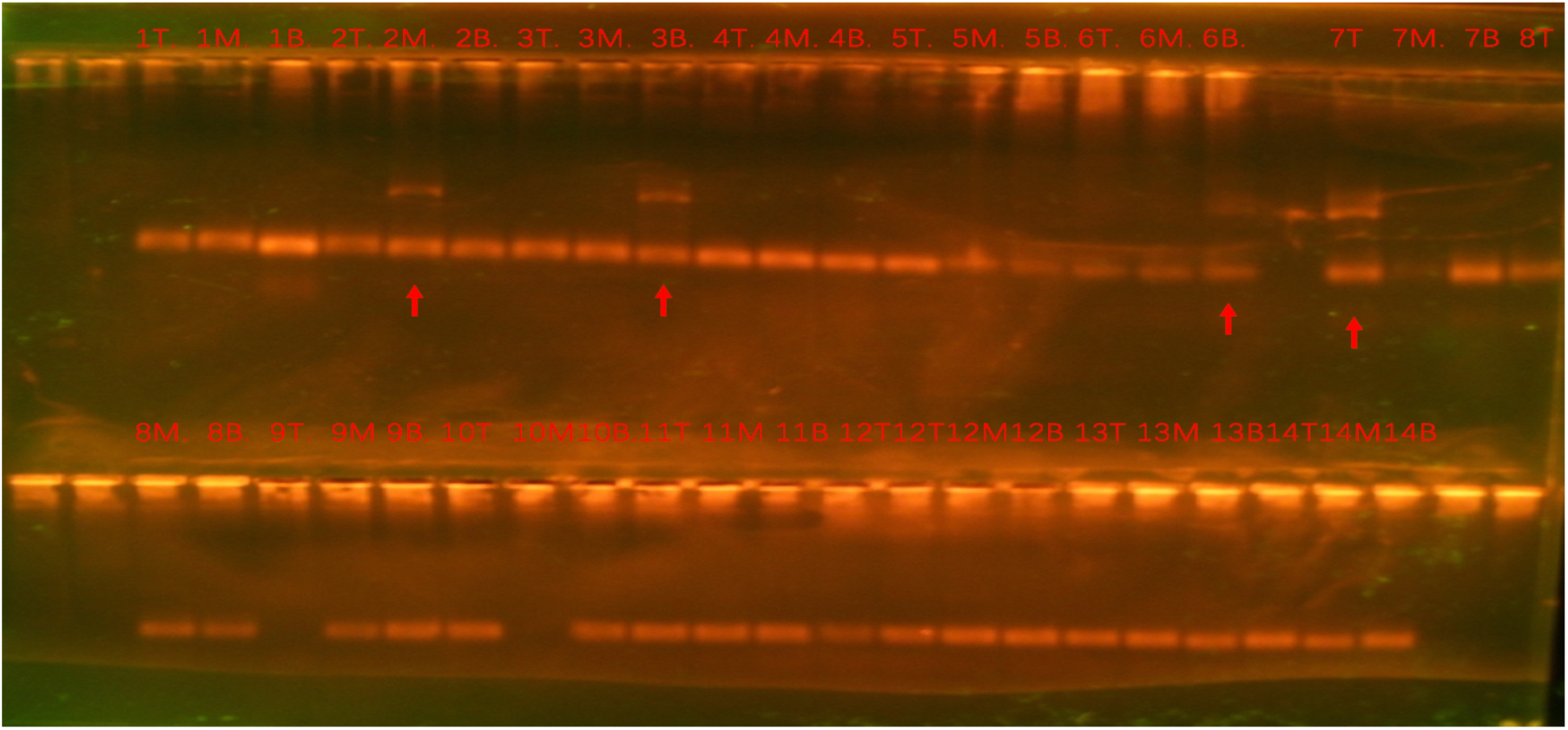
Gel result of 18a/19’ primers. 1.2% agarose gel, 120V, 2 hours. Bio easy marker I and II are used in the left two lanes but didn’t show up. The primer mainly covered the other end of LIS-1 where it joins to original genome. The larger band showed in 2M, 3B,6B,7T

**Figure 9.**
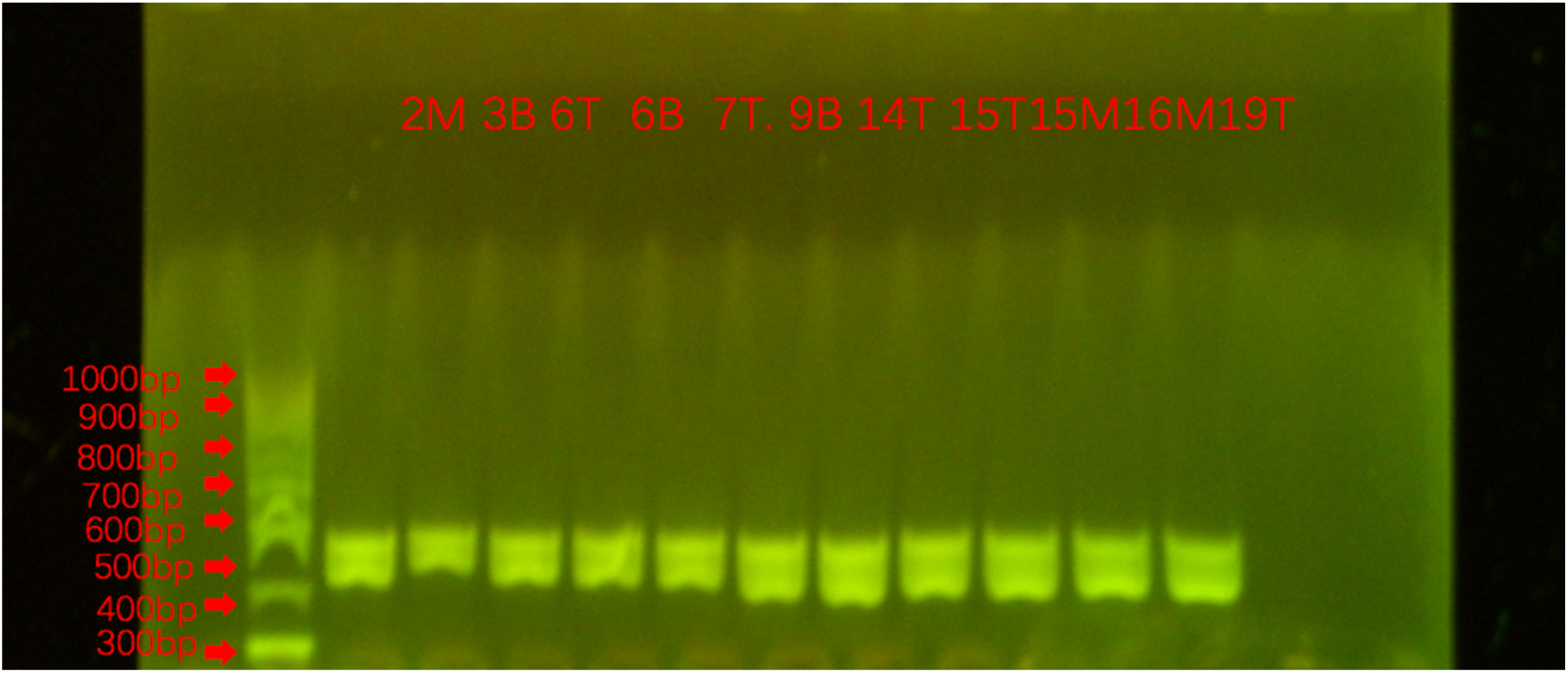
PCR reaction of primer 34. 1.2% agarose gel, 120V, 2 hour. All samples show a wide band covering nearly 100bp range and have slight difference between each other.

IGV suggests a gap at the site in the genome where the primer pair 34 were designed, when comparing the Pl genome with Bethune. The IGV (Figure 10) and previous data (Cullis, 2019) suggests that Pl genotroph has a 500bp band and *Bethune* has an 800bp band in the primer 34 site. All the samples show a Pl character. The wide band may be caused by either high copy number or high original DNA concentration.

**Figure 10.**
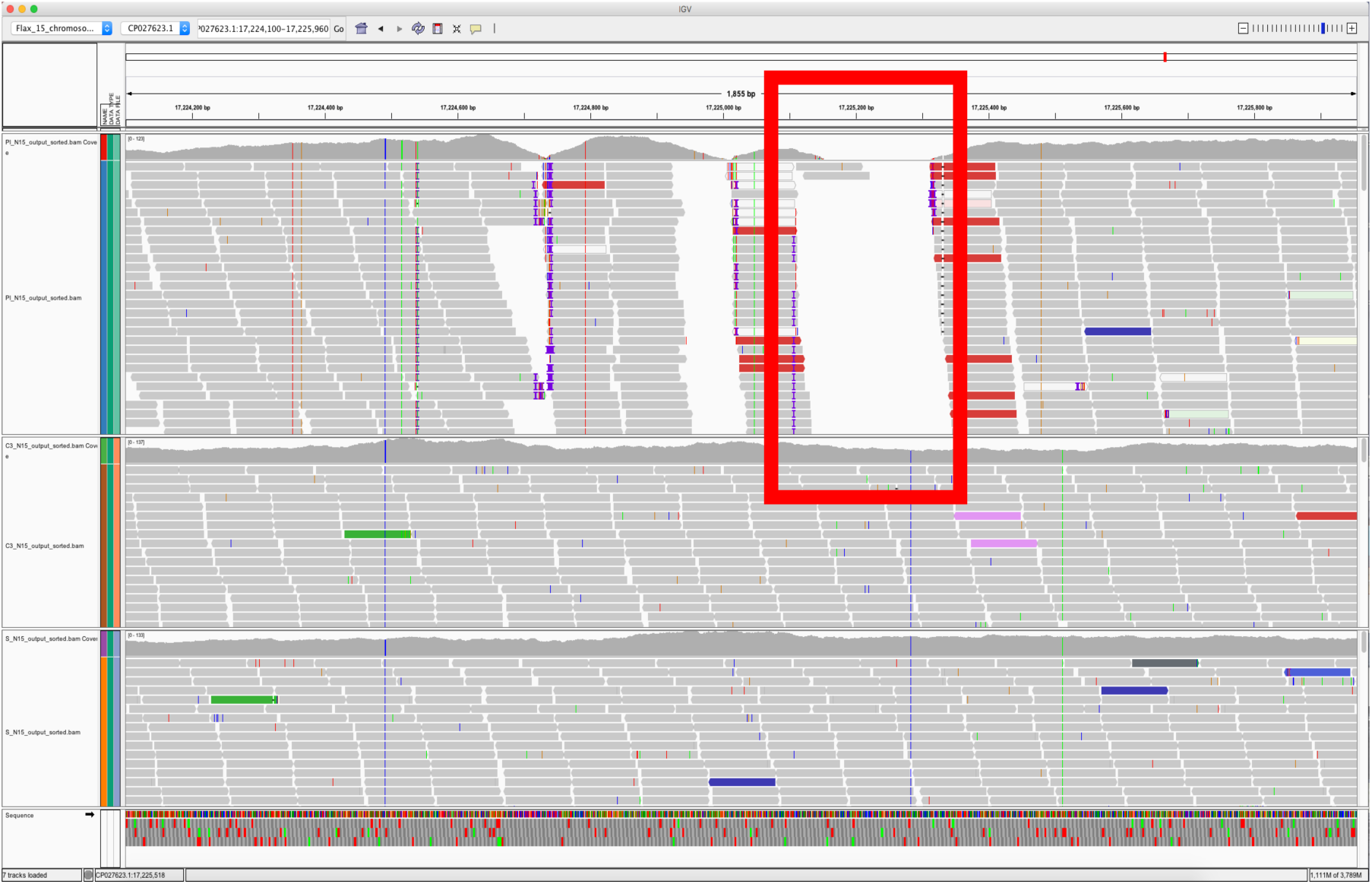
IGV picture of primer 34 site which shows a gap between Pl and C3.

**Figure 11.**
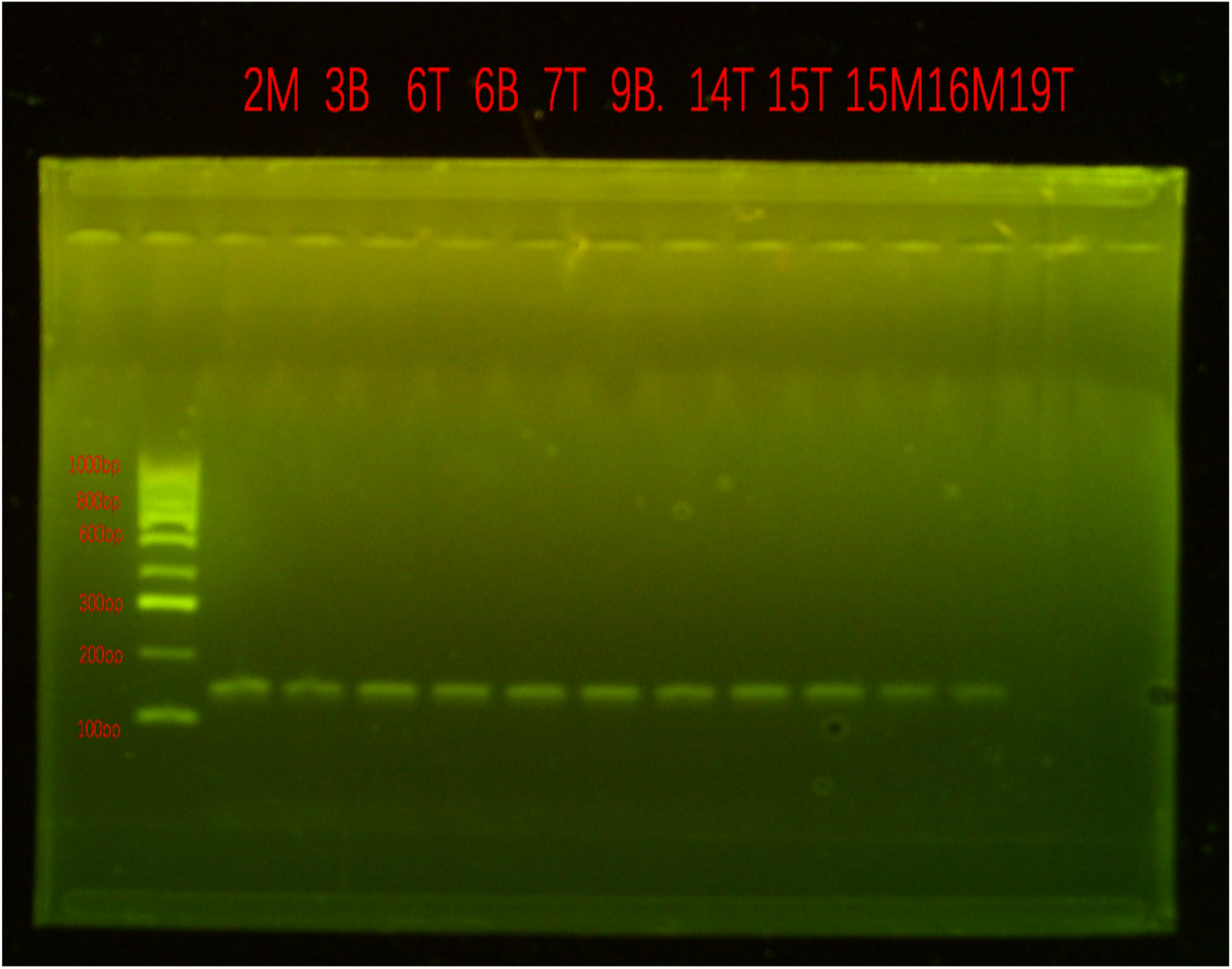
Primer 38. 1.2% agarose gel, 120V, 2 hour. Bio easy marker I and II are used in the left two lanes. Hyperladder IV is used as a marker.

Pl and Bethune show 8bp difference in this site across the region amplified by primer 38 from a previous study (Cullis, 2019). It is not easy to tell the exact size but all samples show a same result.

From table 4, we can see that 2/3’, 2/pl9 and 18a/19’don’t show apparent relationship between each other as previously observed. This means that the LIS-1 is not inserted at one time or is assembled somewhere else before insertion, and the 2/3’ end insertion or 18a/19’ end insertion did not occur at the same time. A similar result was observed previously (Chen, 2005). 4B loses 2/pl9 band when LIS-1 is inserted completely but 2/3’ and 18a/19’ don’t show the expected bands. In terms of the place that the leaves were collected, for 2/3’ primers, 4 top leaves, 3 middle leaves and 2 bottom leaves show extra band; for 2/pl9, 1 bottom sample loses the band; for 18a/19’, 2 top samples, 1 middle samples and 2 bottom samples show extra band. For samples that were selected because of the above variation (2M, 3B, 6B, 6T, 7T, 9B, 14T, 15T, 15M, 16M, 19T), the primer 32 shows all were the same as Pl, but the 32 gap shows the variation that only a small part of the samples containing the band. Primers 34, 38 and 1 all show no variation for the chosen 9 samples and all are the same pattern as Pl. Pl and B are slightly different in the area that primer 33 covered and the samples 9M, 10M, 12T, 13B and 14T have bands same to B/S, but here I cannot rule out the difference caused by the gel due to the small different of the variants. The table 4 shows the percentage of variation happens on different sampling regions, and middle or top samples are more active to the change than bottom leave which suggests a potential development related mechanism. Above all, no obvious linkage between primers was found and variation does show frequency difference due to sampling region. But there is a need for future work to make the conclusion more significant.

**Table 4.**
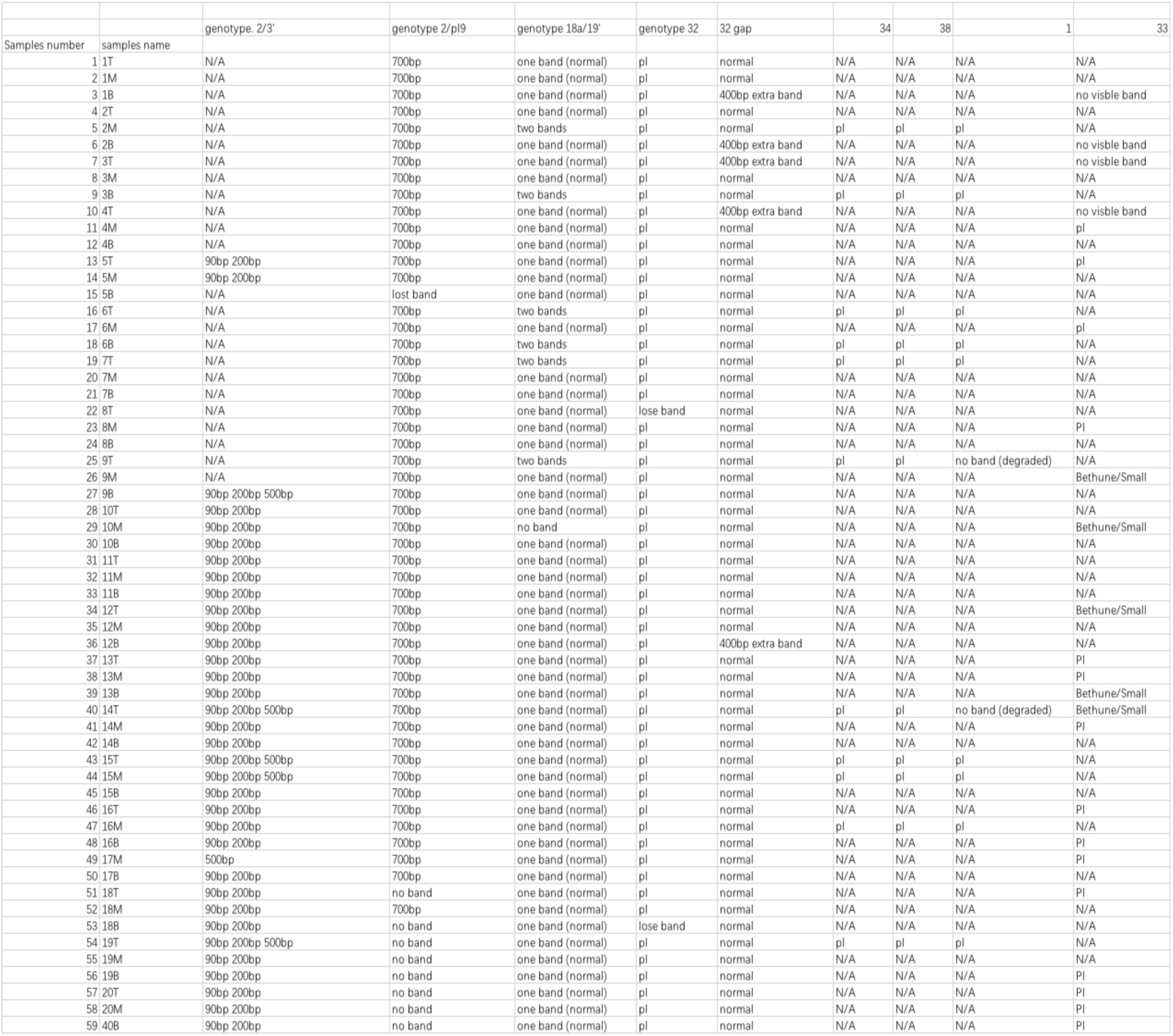
Summary of gel picture result for the first set of samples, Pl refers to Pl band pattern based on Cullis, 2019, extra band refers to the variation bands that different from the others. The blank ones are for faint bands or data is unavailable.

**Table 5.**
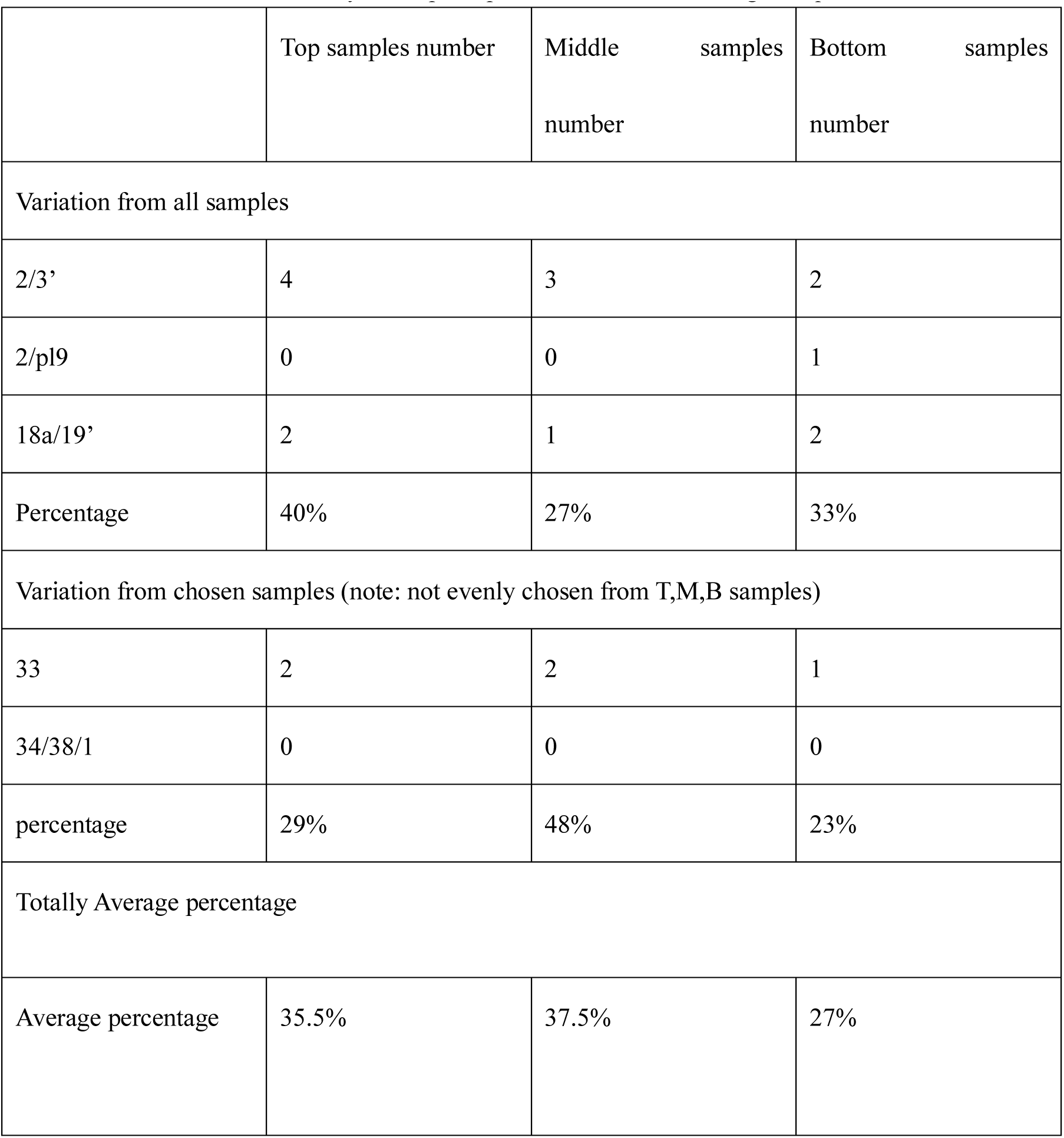
Summary of the place pf an individual for a changed sequence.

**Table 6.**
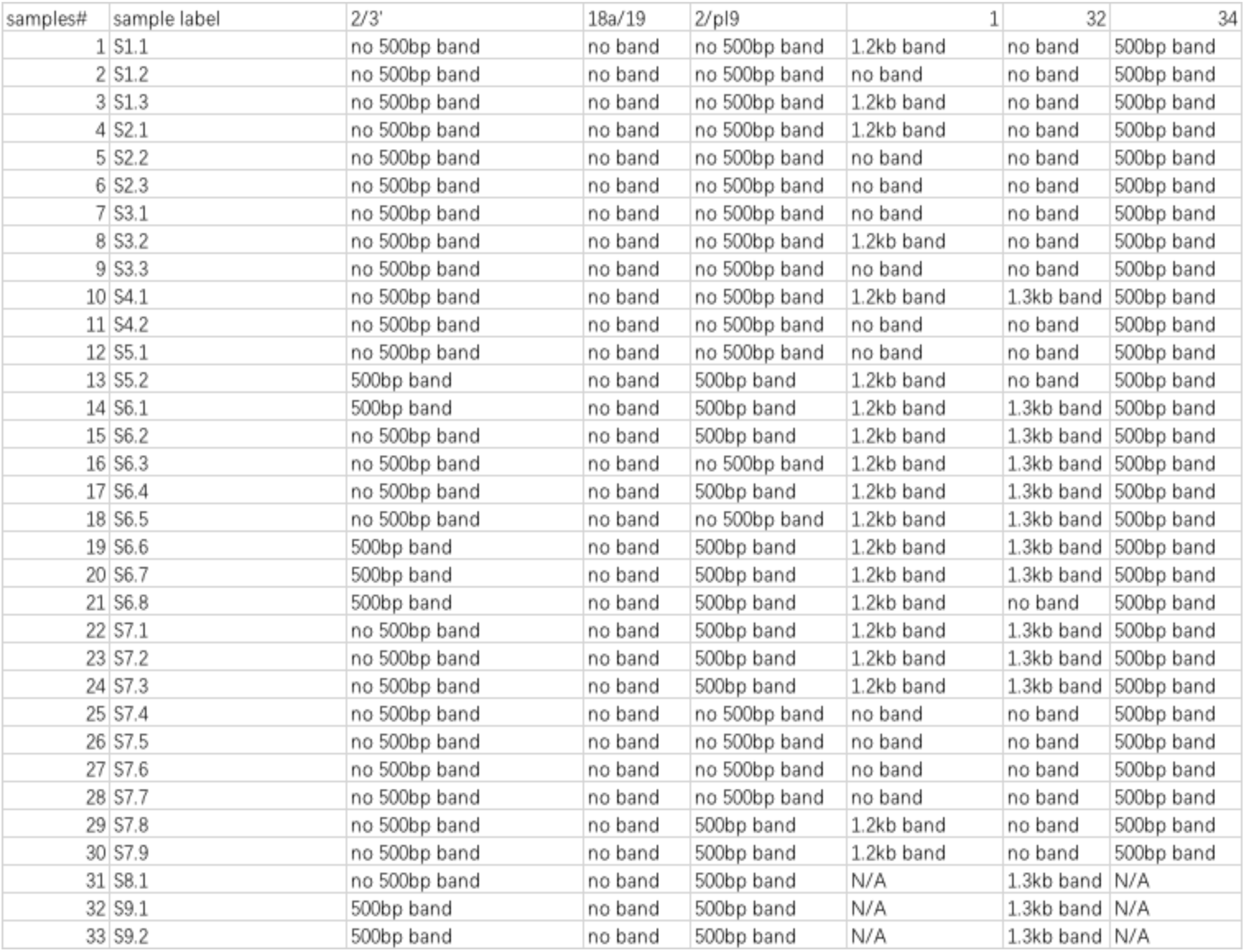
Summary of gel picture result for the second set of samples. The blank means there is no band. N/A means no available data.

| samples# | sample label | 2/3' | 18a/19 | 2/pl9 | 1 | 32 | 34 |
| --- | --- | --- | --- | --- | --- | --- | --- |
| 1 | S1.1 | no 500bp band | no band | no 500bp band | 1.2kb band | no band | 500bp band |
| 2 | S1.2 | no 500bp band | no band | no 500bp band | no band | no band | 500bp band |
| 3 | S1.3 | no 500bp band | no band | no 500bp band | 1.2kb band | no band | 500bp band |
| 4 | S2.1 | no 500bp band | no band | no 500bp band | 1.2kb band | no band | 500bp band |
| 5 | S2.2 | no 500bp band | no band | no 500bp band | no band | no band | 500bp band |
| 6 | S2.3 | no 500bp band | no band | no 500bp band | no band | no band | 500bp band |
| 7 | S3.1 | no 500bp band | no band | no 500bp band | no band | no band | 500bp band |
| 8 | S3.2 | no 500bp band | no band | no 500bp band | 1.2kb band | no band | 500bp band |
| 9 | S3.3 | no 500bp band | no band | no 500bp band | no band | no band | 500bp band |
| 10 | S4.1 | no 500bp band | no band | no 500bp band | 1.2kb band | 1.3kb band | 500bp band |
| 11 | S4.2 | no 500bp band | no band | no 500bp band | no band | no band | 500bp band |
| 12 | S5.1 | no 500bp band | no band | no 500bp band | no band | no band | 500bp band |
| 13 | S5.2 | 500bp band | no band | 500bp band | 1.2kb band | no band | 500bp band |
| 14 | S6.1 | 500bp band | no band | 500bp band | 1.2kb band | 1.3kb band | 500bp band |
| 15 | S6.2 | no 500bp band | no band | 500bp band | 1.2kb band | 1.3kb band | 500bp band |
| 16 | S6.3 | no 500bp band | no band | no 500bp band | 1.2kb band | 1.3kb band | 500bp band |
| 17 | S6.4 | no 500bp band | no band | 500bp band | 1.2kb band | 1.3kb band | 500bp band |
| 18 | S6.5 | no 500bp band | no band | no 500bp band | 1.2kb band | 1.3kb band | 500bp band |
| 19 | S6.6 | 500bp band | no band | 500bp band | 1.2kb band | 1.3kb band | 500bp band |
| 20 | S6.7 | 500bp band | no band | 500bp band | 1.2kb band | 1.3kb band | 500bp band |
| 21 | S6.8 | 500bp band | no band | 500bp band | 1.2kb band | no band | 500bp band |
| 22 | S7.1 | no 500bp band | no band | 500bp band | 1.2kb band | 1.3kb band | 500bp band |
| 23 | S7.2 | no 500bp band | no band | 500bp band | 1.2kb band | 1.3kb band | 500bp band |
| 24 | S7.3 | no 500bp band | no band | 500bp band | 1.2kb band | 1.3kb band | 500bp band |
| 25 | S7.4 | no 500bp band | no band | no 500bp band | no band | no band | 500bp band |
| 26 | S7.5 | no 500bp band | no band | no 500bp band | no band | no band | 500bp band |
| 27 | S7.6 | no 500bp band | no band | no 500bp band | no band | no band | 500bp band |
| 28 | S7.7 | no 500bp band | no band | no 500bp band | no band | no band | 500bp band |
| 29 | S7.8 | no 500bp band | no band | 500bp band | 1.2kb band | no band | 500bp band |
| 30 | S7.9 | no 500bp band | no band | 500bp band | 1.2kb band | no band | 500bp band |
| 31 | S8.1 | no 500bp band | no band | 500bp band | N/A | 1.3kb band | N/A |
| 32 | S9.1 | 500bp band | no band | 500bp band | N/A | 1.3kb band | N/A |
| 33 | S9.2 | 500bp band | no band | 500bp band | N/A | 1.3kb band | N/A |

**Table 7.** The number of the sample means different individual, while the “w” refers to water, n: nutrition; l: leaves; m: meristem. The first thirteen samples are collected on date? With a 5-week growth, while the red- and-strengthened named samples are collected 2 weeks after the first set (7-8 weeks growth time).

| Samples | 2/3' | 2/pl9 | 18a/19' | 1 | 2 | 6 | 34 |
| --- | --- | --- | --- | --- | --- | --- | --- |
| 1wl | 90bp band and 200bp band | 500bp band | no band | 800bp band | 400bp band | 600bp band | 480bp band |
| 1nl | 90bp band and 200bp band | 500bp band | no band | 800bp band | 400bp band | 600bp band | 480bp band |
| 2wl | 90bp band and 200bp band and unexpected 800bp band | 500bp band | no band | **200bp band** | **450bp band** | **900bp band** | 480bp band |
| 2wm | 90bp band and 200bp band | 500bp band | no band | **200bp band** | **450bp band** | **900bp band** | 480bp band |
| 2nl | 90bp band and 200bp band | 500bp band | no band | 800bp band | 400bp band | 600bp band | 480bp band |
| 2nm | 90bp band and 200bp band | 500bp band | no band | 800bp band | 400bp band | 600bp band | 480bp band |
| 3wl | 90bp band and 200bp band | 500bp band | no band | 800bp band | 400bp band | 600bp band | 480bp band |
| 3wm | 90bp band and 200bp band | 500bp band | no band | 800bp band | 400bp band | 600bp band | 480bp band |
| 3nl | 90bp band and 200bp band | 500bp band | no band | 800bp band | 400bp band | 600bp band | 480bp band |
| 3nm | 90bp band and 200bp band | 500bp band | no band | 800bp band | 400bp band | 600bp band | 480bp band |
| 4wl | 90bp band and 200bp band | 500bp band | no band | 800bp band | 400bp band | 600bp band | 480bp band |
| 4wm | 90bp band and 200bp band | 500bp band | no band | 800bp band | 400bp band | 600bp band | 480bp band |
| 4nl | 90bp band and 200bp band | 500bp band | no band | 800bp band | 400bp band | 600bp band | 480bp band |
| **3wl** | 90bp band and 200bp band | 500bp band | no band | 800bp band | 400bp band | 600bp band | 480bp band |
| **3wm** | 90bp band and 200bp band | 500bp band | no band | 800bp band | 400bp band | 600bp band | 480bp band |
| **3nl** | 90bp band and 200bp band | 500bp band | no band | 800bp band | 400bp band | 600bp band | 480bp band |
| **3nm** | 90bp band and 200bp band | 500bp band | no band | 800bp band | 400bp band | 600bp band | 480bp band |
| **4wl** | 90bp band and 200bp band | 500bp band | no band | 800bp band | 400bp band | 600bp band | 480bp band |
| **4wm** | 90bp band and 200bp band | 500bp band | no band | 800bp band | 400bp band | 600bp band | 480bp band |
| **4nl** | 90bp band and 200bp band | 500bp band | no band | 800bp band | 400bp band | 600bp band | 480bp band |
| **4nm** | 90bp band and 200bp band | 500bp band | no band | 800bp band | 400bp band | 600bp band | 480bp band |

### 2: Samples collected from different branches of the individuals

DNA was extracted from different branches of individuals. All the individuals are Pl grown in low nutrient environment. The sample were labelled with an “S” before sample number, however, this “S” just refers to “Samples” but is not related in any way to the genotroph S.

Figure 12 refers to result of DNAs amplified by primer 32. The previous data provide two band sizes with this primer pair, one is around 1kb band, anther band is around 5 kb. The 5 kb band would be expected to show up for the samples having no band, but the elongation time used for these reactions was insufficient to amplify the 5kb band. The presence of the 5kb band needs to be demonstrated for those samples which gave no amplification.

**Figure 12.**
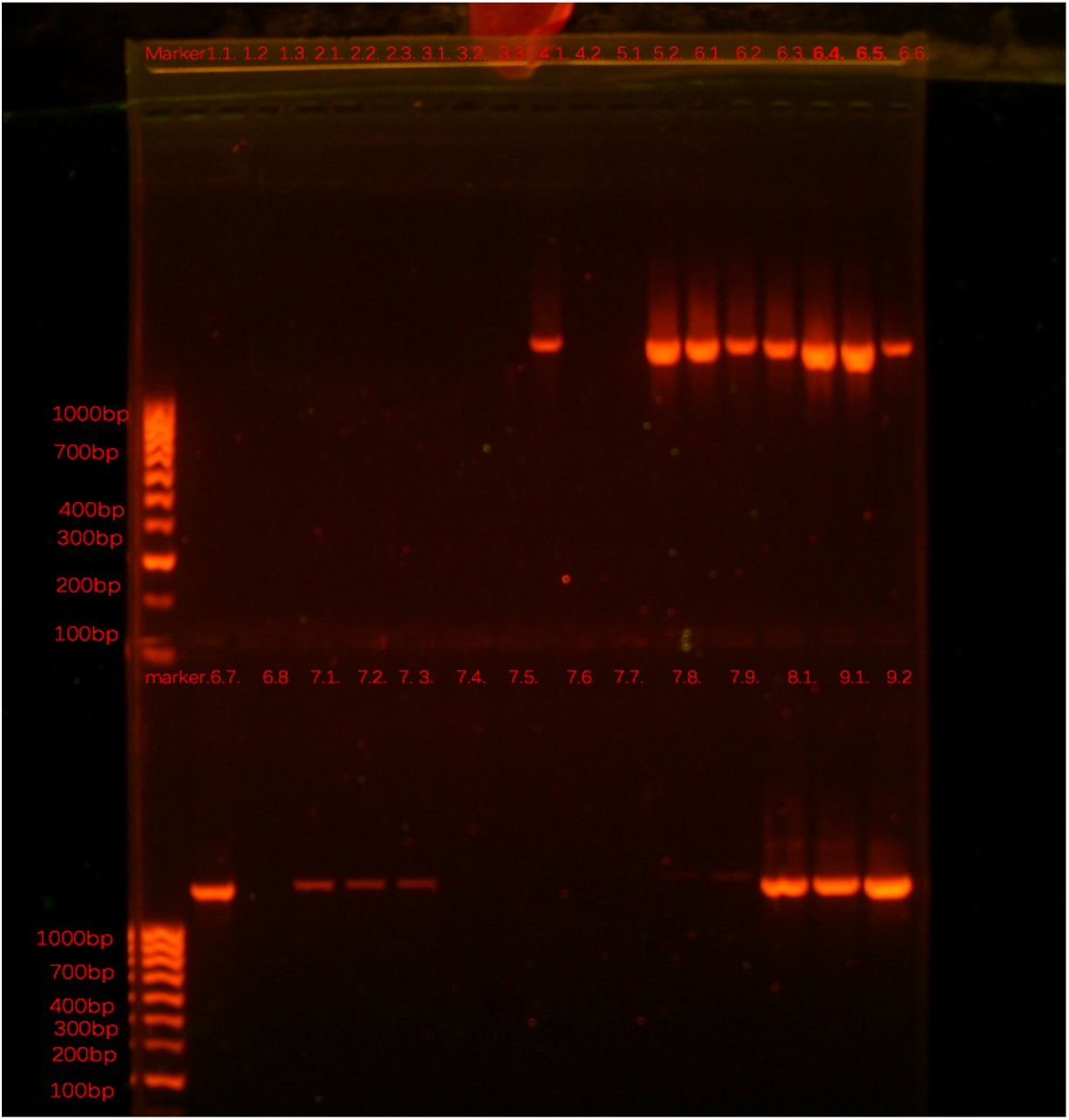
PCR reaction with Primers 32. Some samples show bands over 1kb, while the other lose the band.

**Figure 13.**
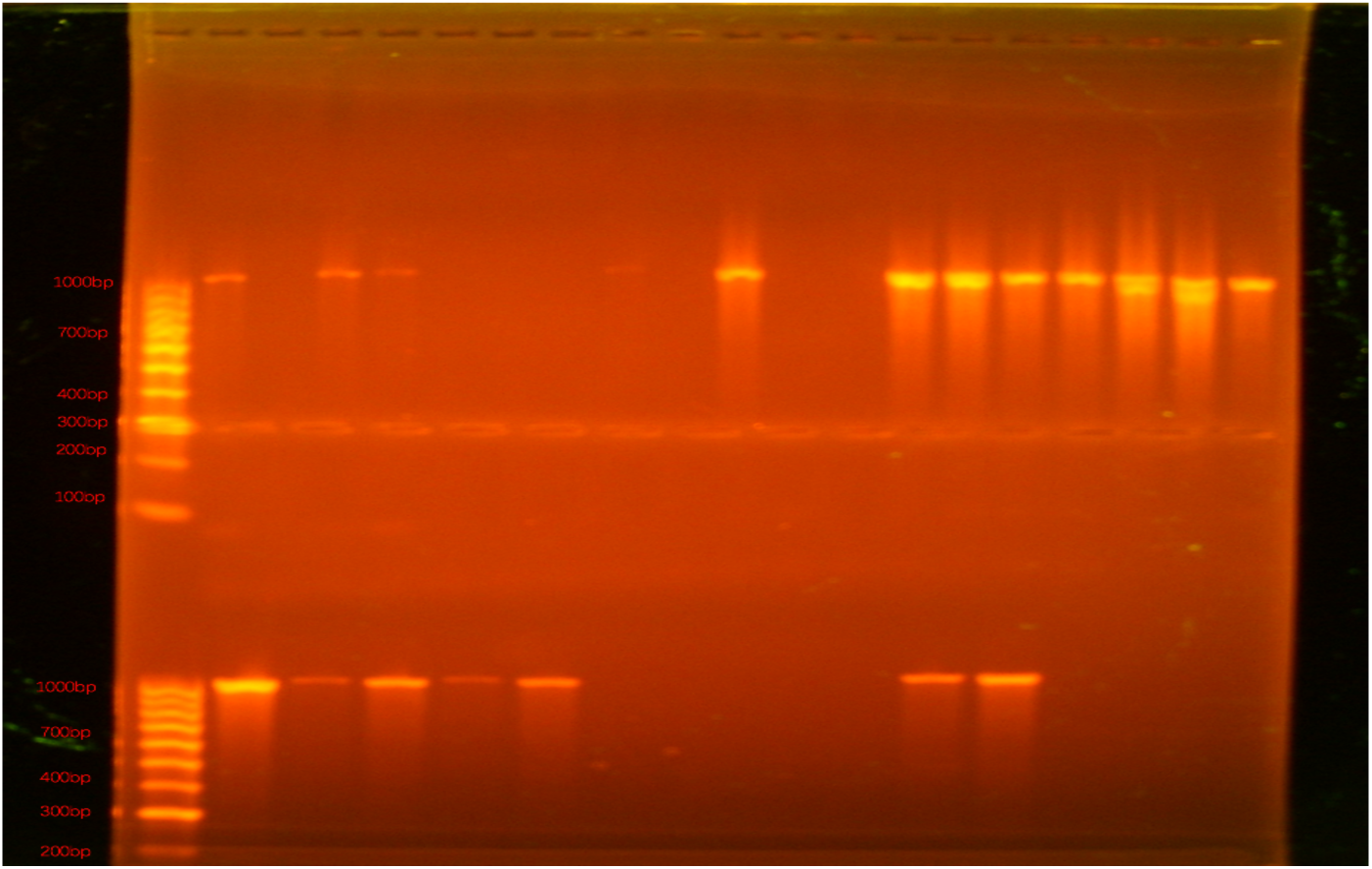
PCR amplifying with primer 1. The gel order :1:Marker (hyperladder iv) 2:S1.1. 3:S1.2 4:S1.3. 5:S2.1. 6:S2.2. 7:S2.3. 8:S3.1. 9:S3.2. 10:S3.3. 11:S4.1. 12:S4.2. 13:S5.1 14:S5.2. 15:S6.1. 16:S6.2. 17:S6.3. 18:S6.4. 19:S6.5. 20:S6.6. 21:marker. 22:S6.7. 23: S6.8 24:S7.1. 25:S7.2. 26:S7.3 27:S7.4. 28:S7.5. 29: S7.6 30:S 7.7. 31:S7.8. 32:S7.9. Gel condition: 120V, 1% agarose gel for 4 hours

**Figure 14.**
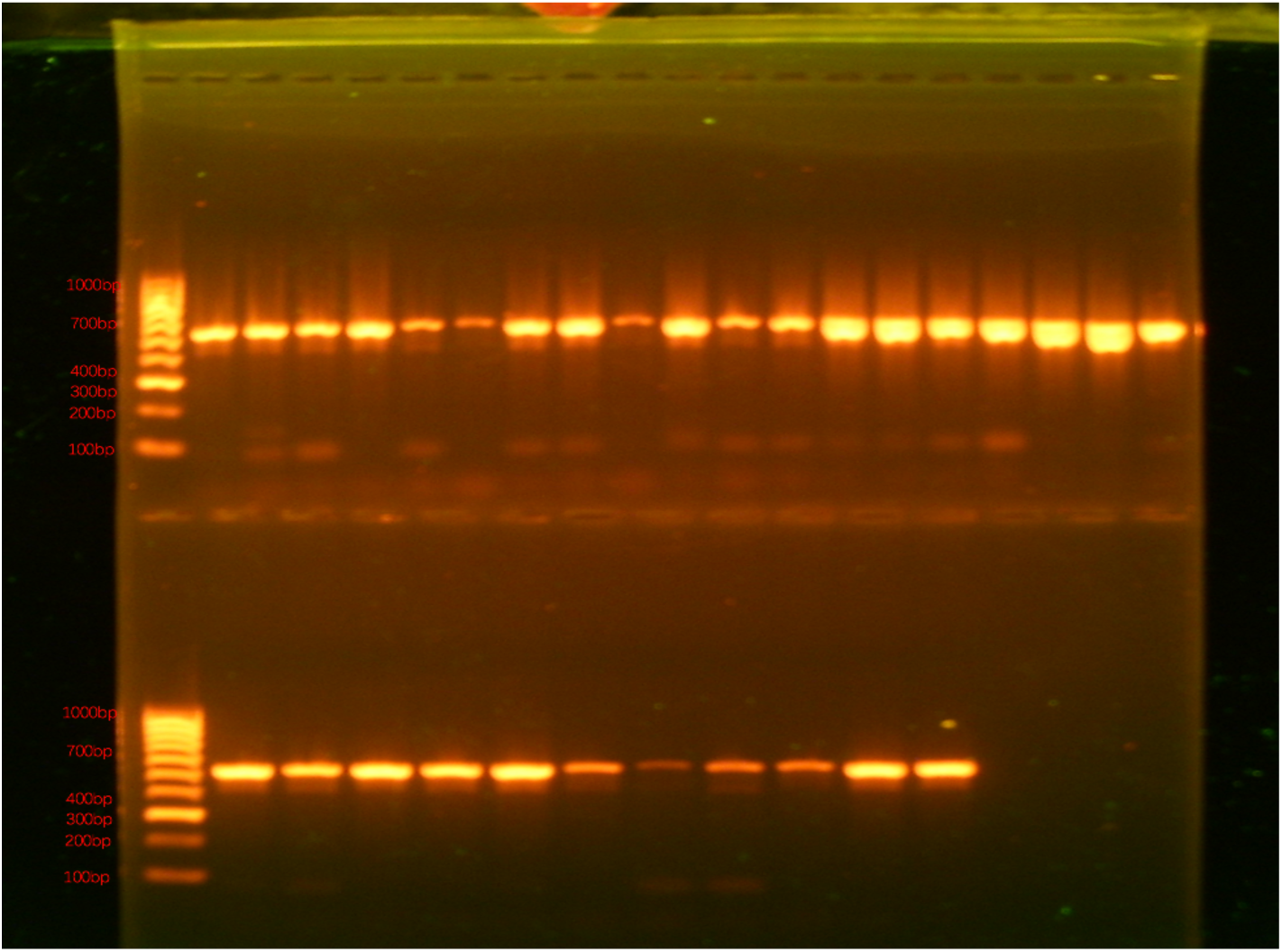
PCR amplifying with primer 34. The gel order :1:Marker (hyperladder iv) 2:S1.1. 3:S1.2 4:S1.3. 5:S2.1. 6:S2.2. 7:S2.3. 8:S3.1. 9:S3.2. 10:S3.3. 11:S4.1. 12:S4.2. 13:S5.1 14:S5.2. 15:S6.1. 16:S6.2. 17:S6.3. 18:S6.4. 19:S6.5. 20:S6.6. 21:marker. 22:S6.7. 23: S6.8 24:S7.1. 25:S7.2. 26:S7.3 27:S7.4. 28:S7.5. 29: S7.6 30:S 7.7. 31:S7.8. 32:S7.9. Gel condition: 120V, 1% agarose gel for 4 hours

**Figure 15.**
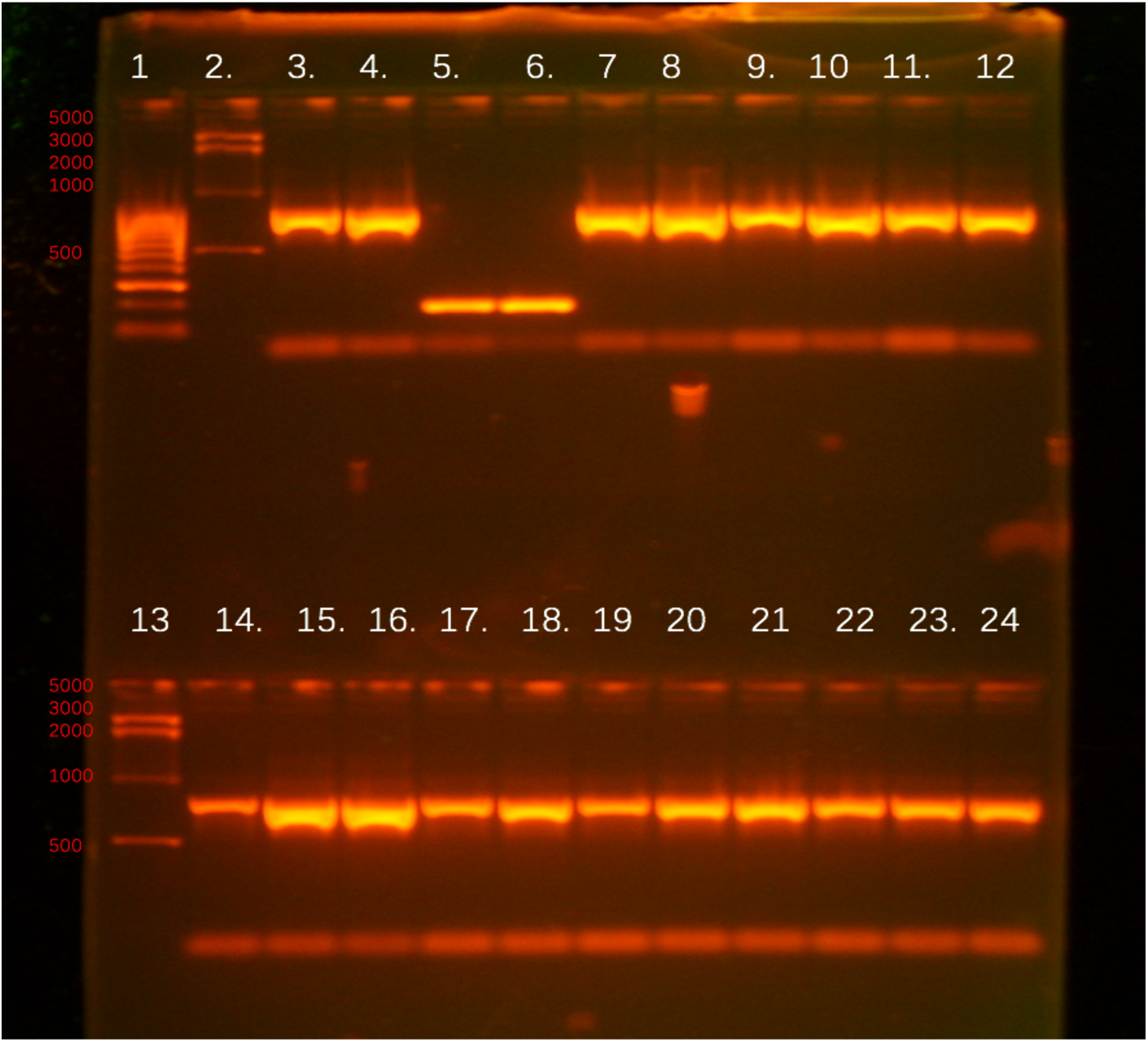
PCR reaction with primer 1. Order 1: marker 1 2: marker 2 3:1wl. 4:1nl 5:2wl 6:2wm 7:2nl. 8:2nm 9:3wl 10:2wm 11:3nl 12:3nm 14:4wl 15:4wm 16:4nl 17:3wl 18:3wm 19:3nl 20:3nm 21:4wl 22:4wm 23:4nl 24:4nm. Gel condition: 1.2%, 100V, 2hours.

For this set of plants, the 18a/19 primer shows no band for all samples, however, 7 samples have 2/3’band, indicating that part of the LIS-1 insertion is formed from 2/3’ end. All samples having 2/3’band also have the 2/pl9 band but not all samples having 2/pl9 band show 2/3’ ones that may indicate that the 3’ sequence not has to be inserted first specifically. For primer 1, there are also some samples losing the band. This is consistent with the statement that the primer 1 covers a transposon element site which is around 1kb length (Wilkinson personal communication, 2019). Based on the gap, the transposon could be a cut and paste type. The footprint of this deletion reinforced the potential transposon nature of primer 1-covered sequence (Figure 19). After alignment of this sequence in BLASTn, there are two matches with repeat sequence in my sequence (Figure 19). After amplification with primer 32, some samples show 1.1kb bands, which is consistent with previous result that 32 primer should cover a site with variation of 1kb and 5kb. The 5kb band is not amplified successfully in this experiment because of the elongation time is long not enough for the PCR quality. For primer 34, all samples have a 500bp band which is consistent with the Pl character.

**Figure 16.**
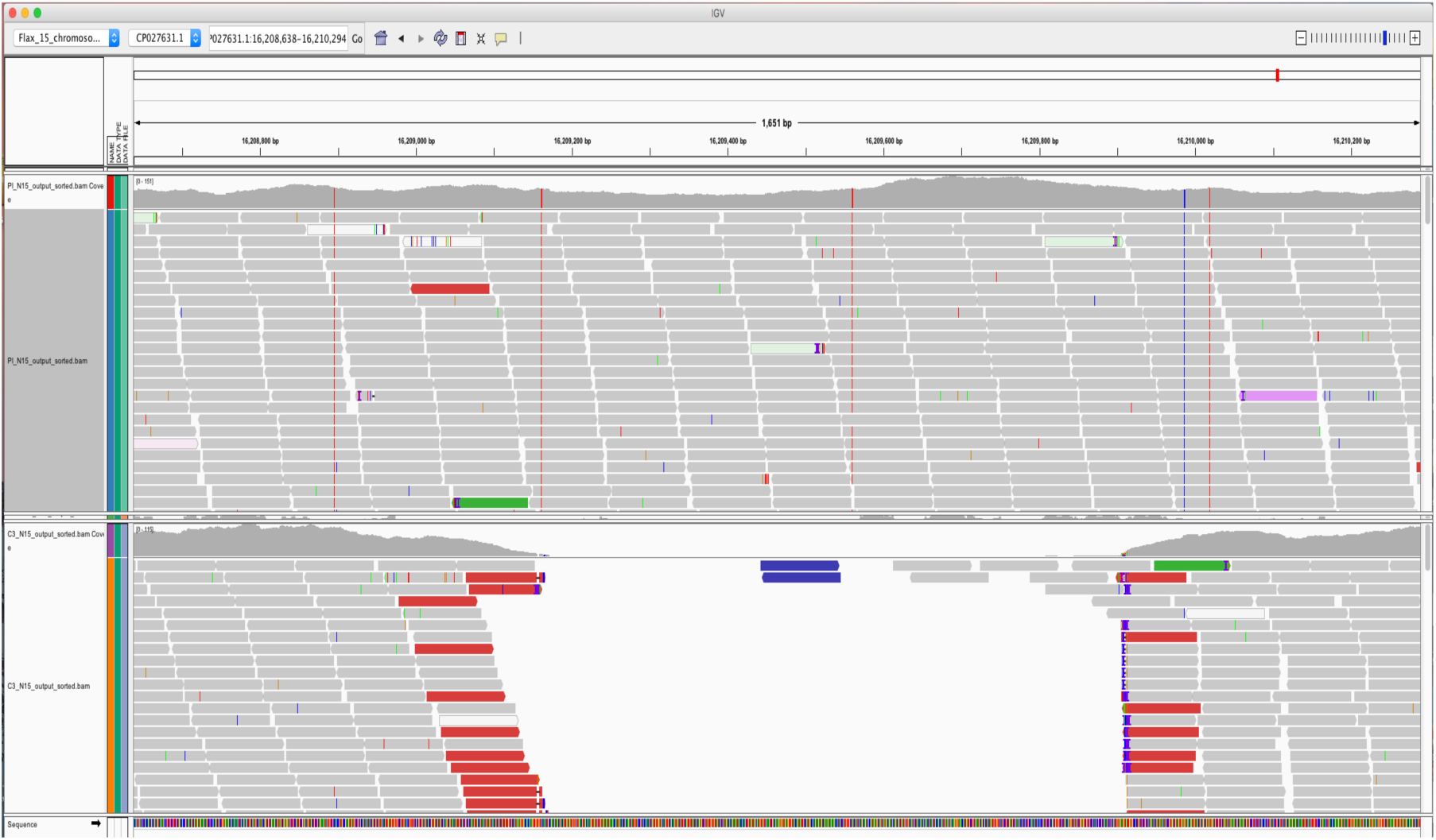
IGV picture at site 1. The Bethune is used as a reference sequence. Compared Pl to C3, there are two obvious SNPs. In Pl, there is also a gap around 800bp between 16209200 to 16209900 sequencing result.

**Figure 17.**
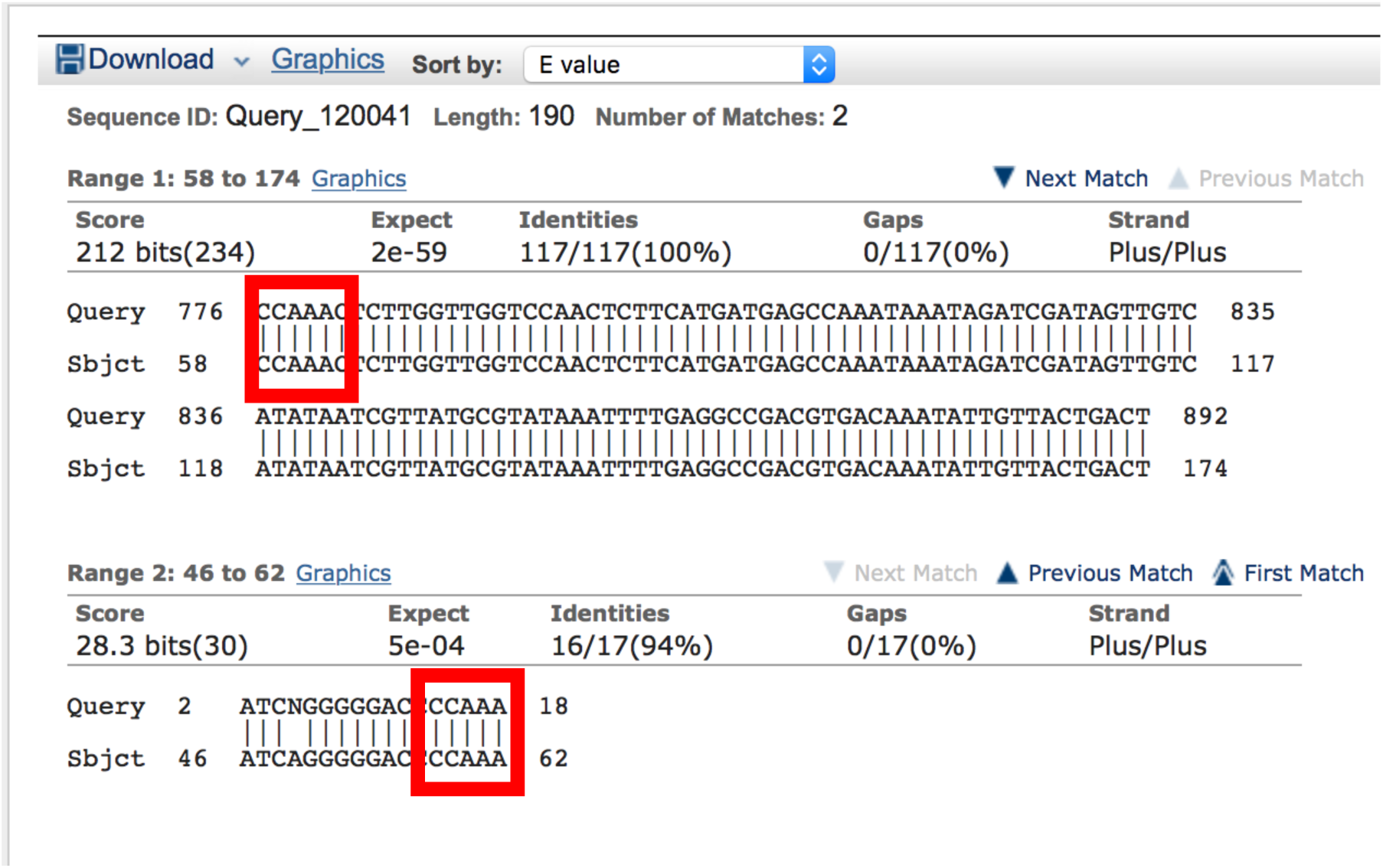
BLAST result of PCR reaction amplified by primer 1. Two samples that show difference between each other are chosen.

**Figure 18.**
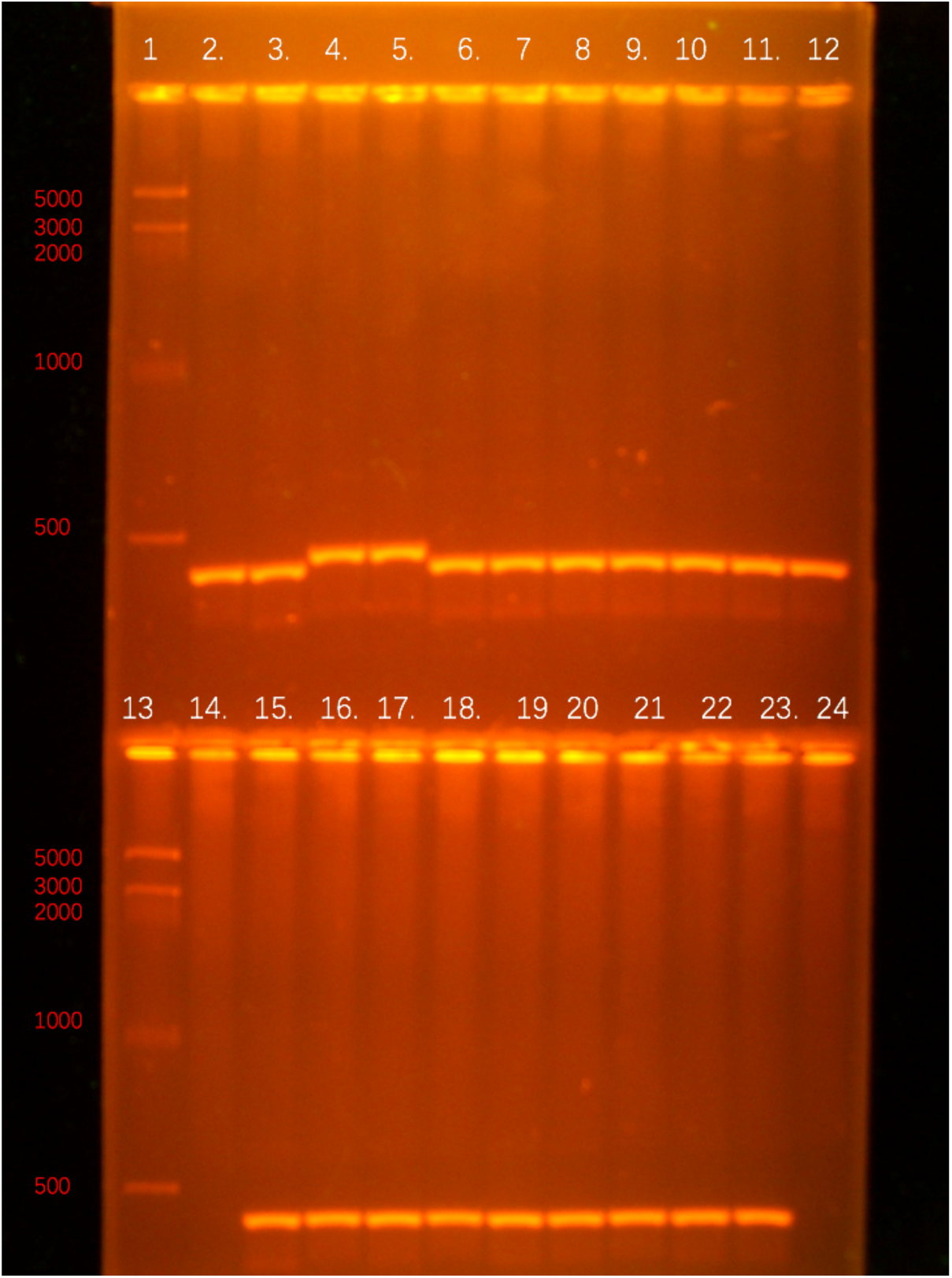
PCR reaction with primer 2. Order: 1: marker 1 2: marker 2 3:1wl. 4:1nl 5:2wl 6:2wm 7:2nl. 8:2nm 9:3wl 10:2wm 11:3nl 12:3nm 14:4wl 15:4wm 16:4nl 17:3wl 18:3wm 19:3nl 20:3nm 21:4wl 22:4wm 23:4nl 24:4nm. Gel condition: 1.2%, 100V, 2hours.

**Figure 19.**
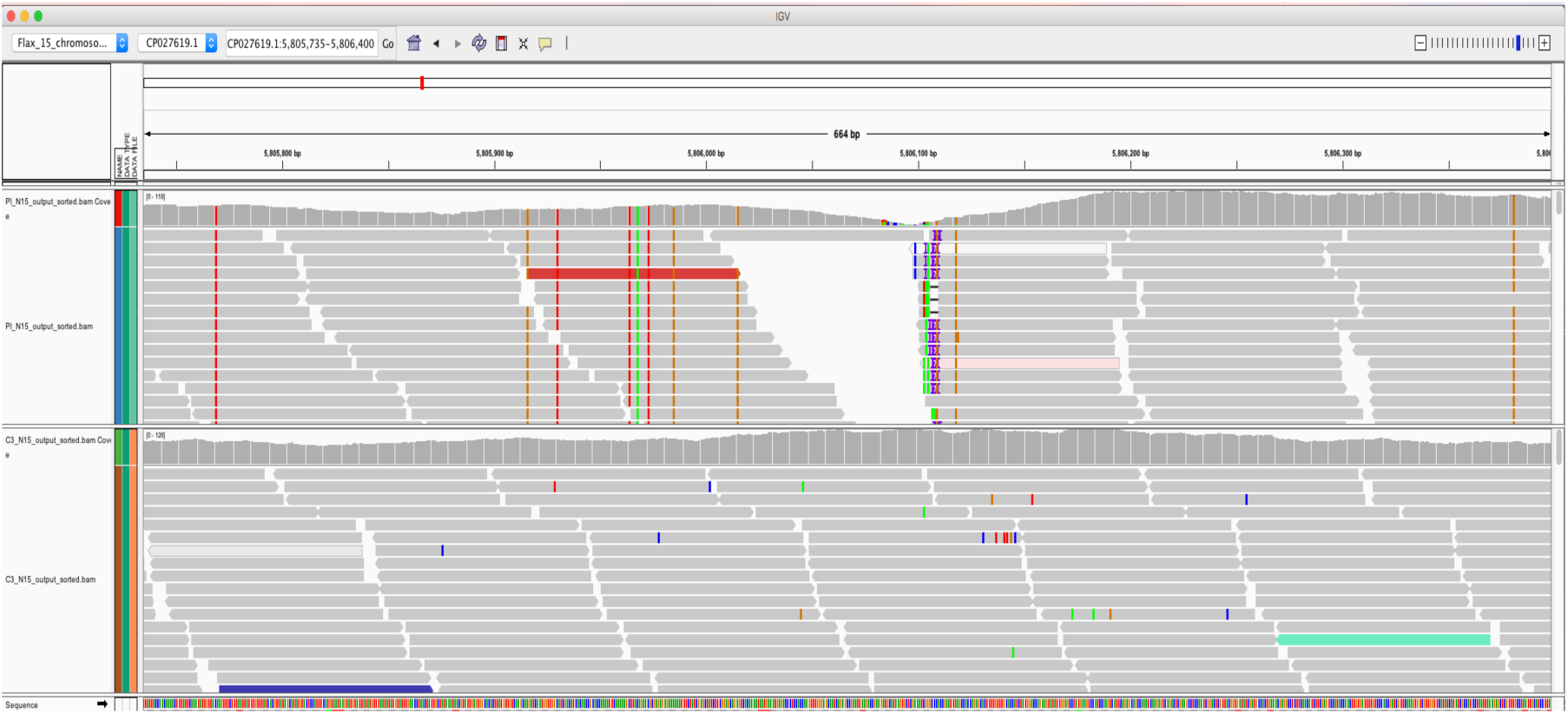
IGV picture at site 2. There is a small gap (around 10bp) between Pl and Bethune, and also some SNPs around this site. However, the C3, which is derived from Pl, becomes more similar to Bethune.

**Figure 20.**
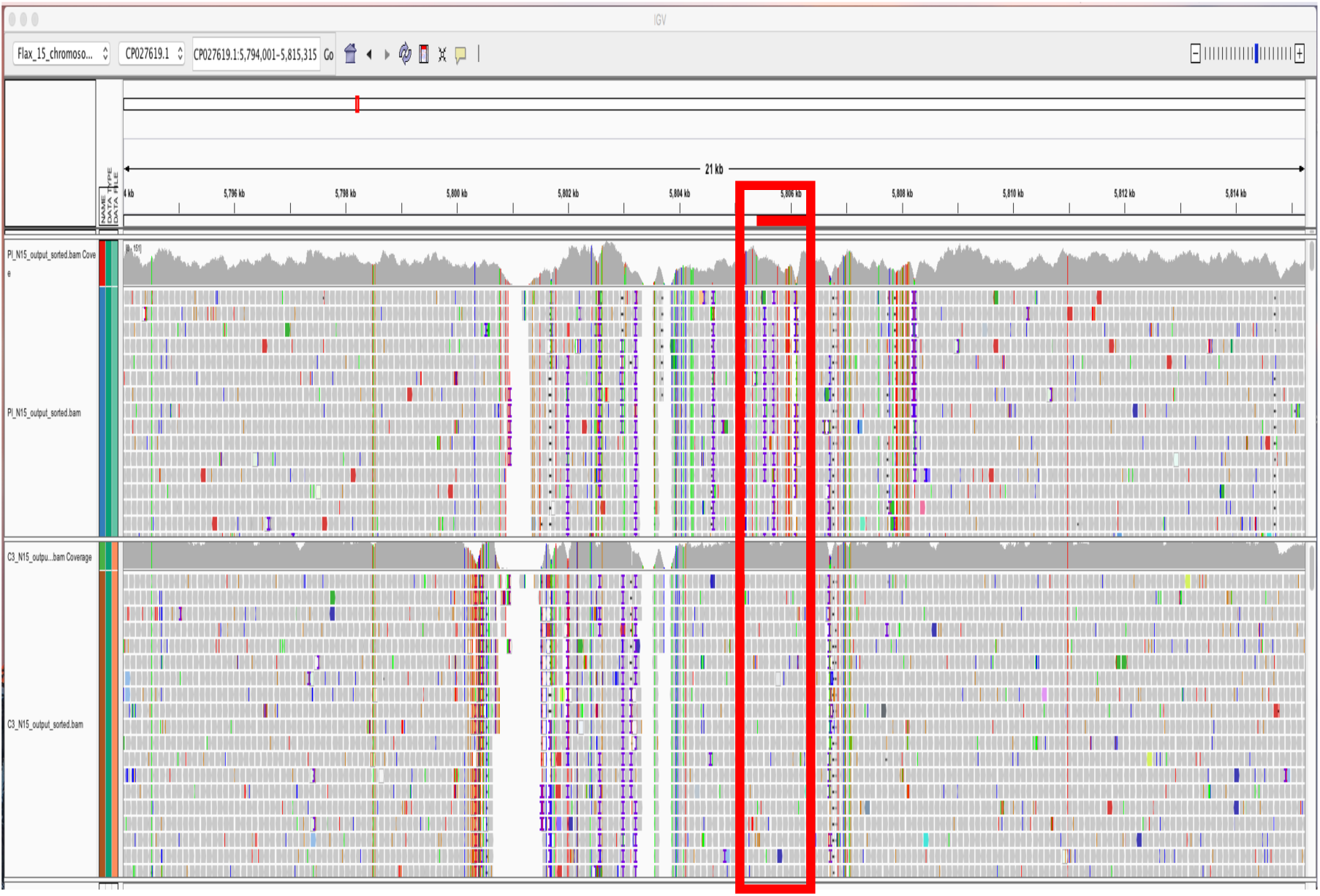
IGV picture at primer site 2 but the picture contains more information about the region of primer 2, where many polymorphisms come up. The square means the primer 2 site.

**Figure 21.**
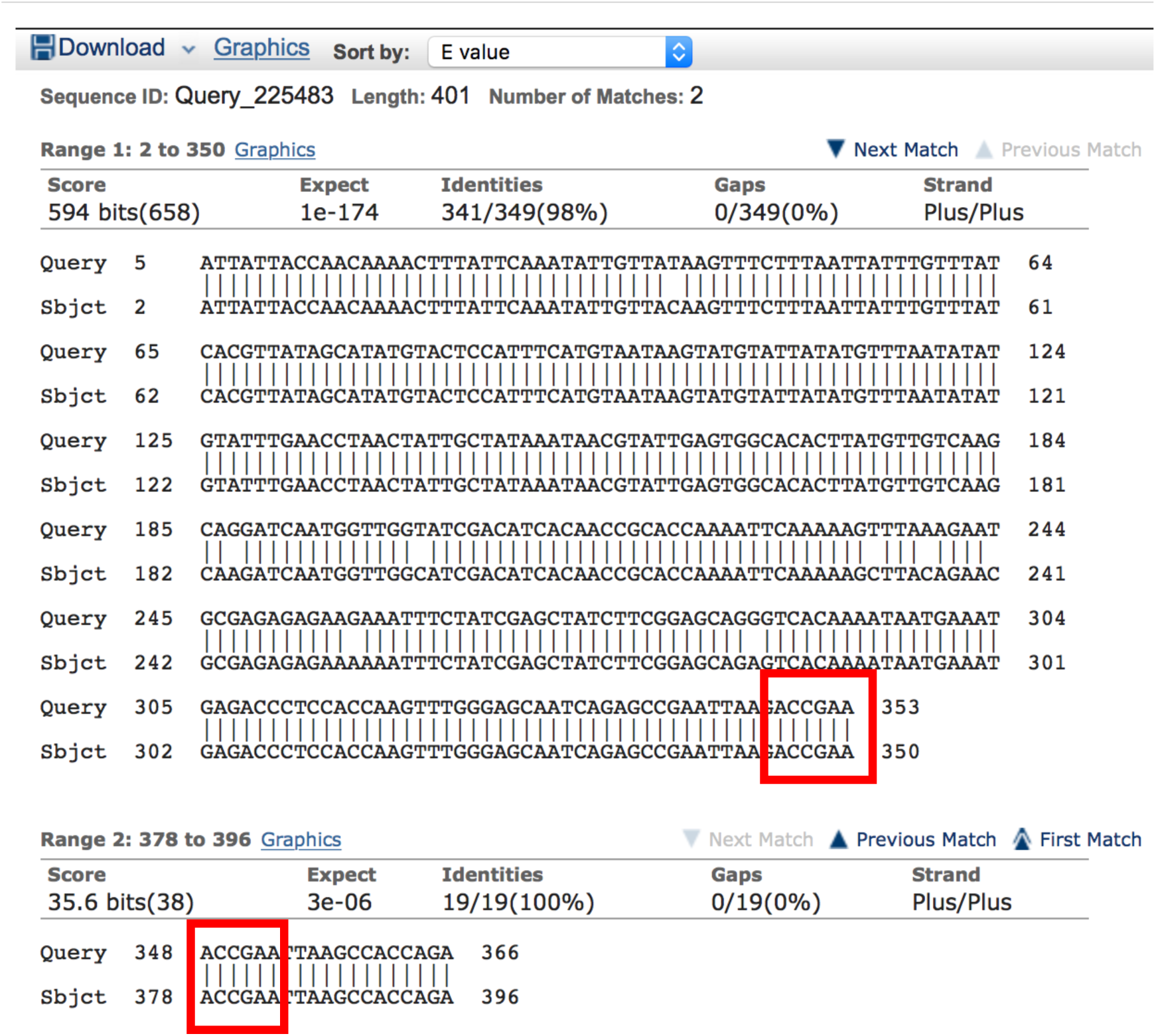
BLAST result of PCR reaction amplified by primer 2 of two samples (1nl and 2wl) that show difference between each other on a gel.

**Figure 22.**
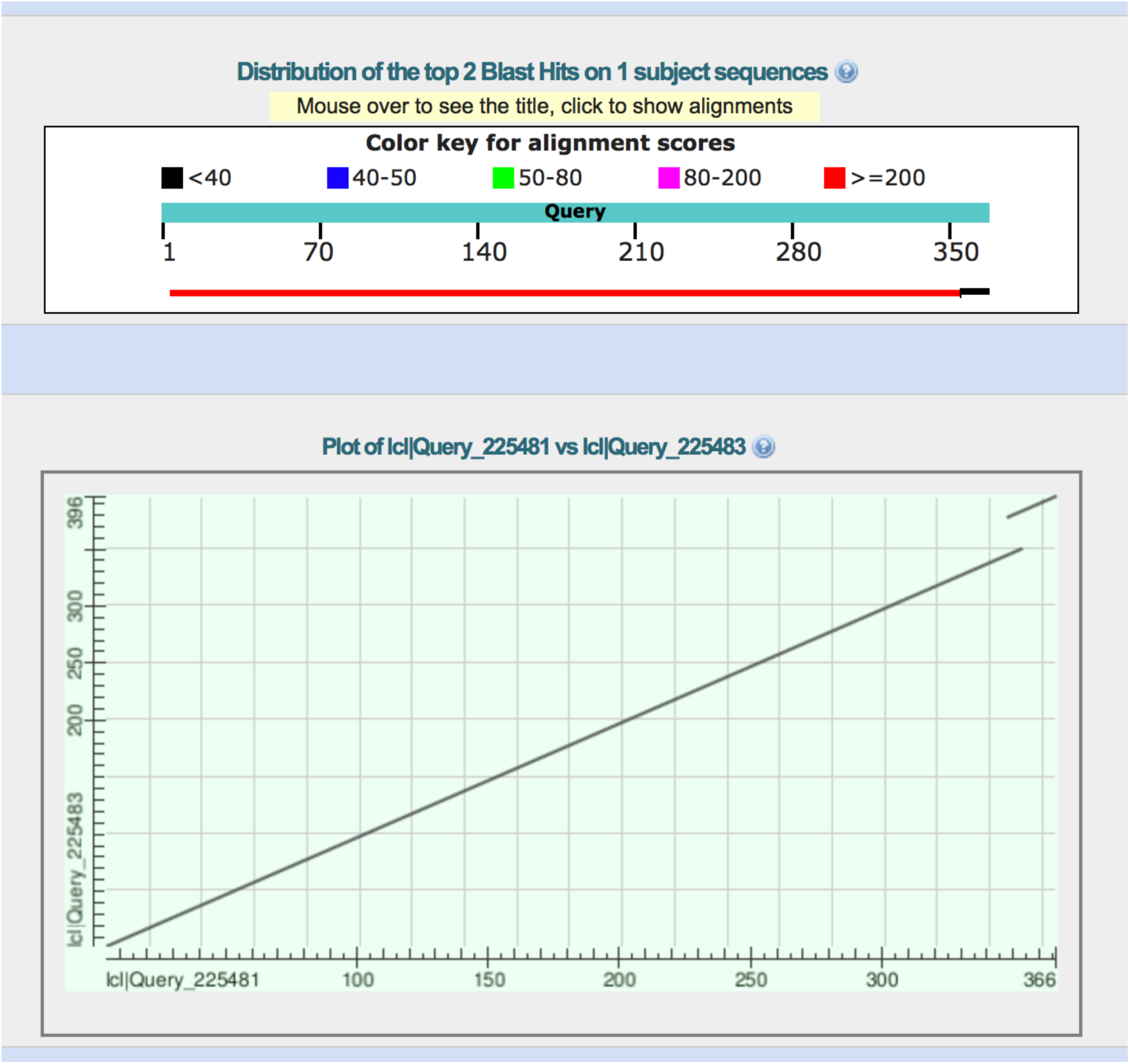
Plot picture of primer 2 Blast result.

**Figure 23.**
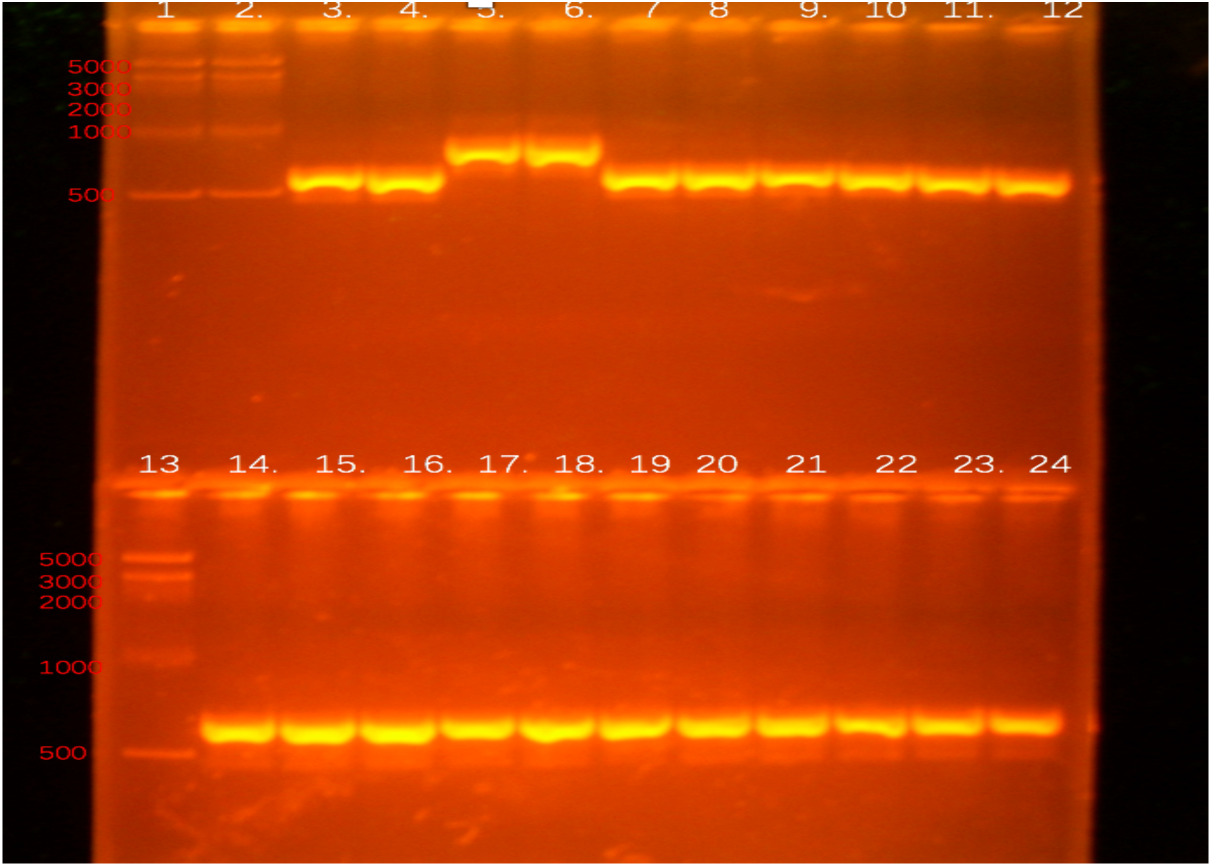
PCR amplification with primer 6. Order of samples. 1: marker II 2:marker II 3:1wl. 4:1nl 5:2wl 6:2wm 7:2nl. 8:2nm 9:3wl 10:2wm 11:3nl 12:3nm 13: marker II 14:4wl 15:4wm 16:4nl 17:3wl 18:3wm 19:3nl 20:3nm 21:4wl 22:4wm 23:4nl 24:4nm. Gel condition: 100V, 1.2%, 2 hours

**Figure 24.**
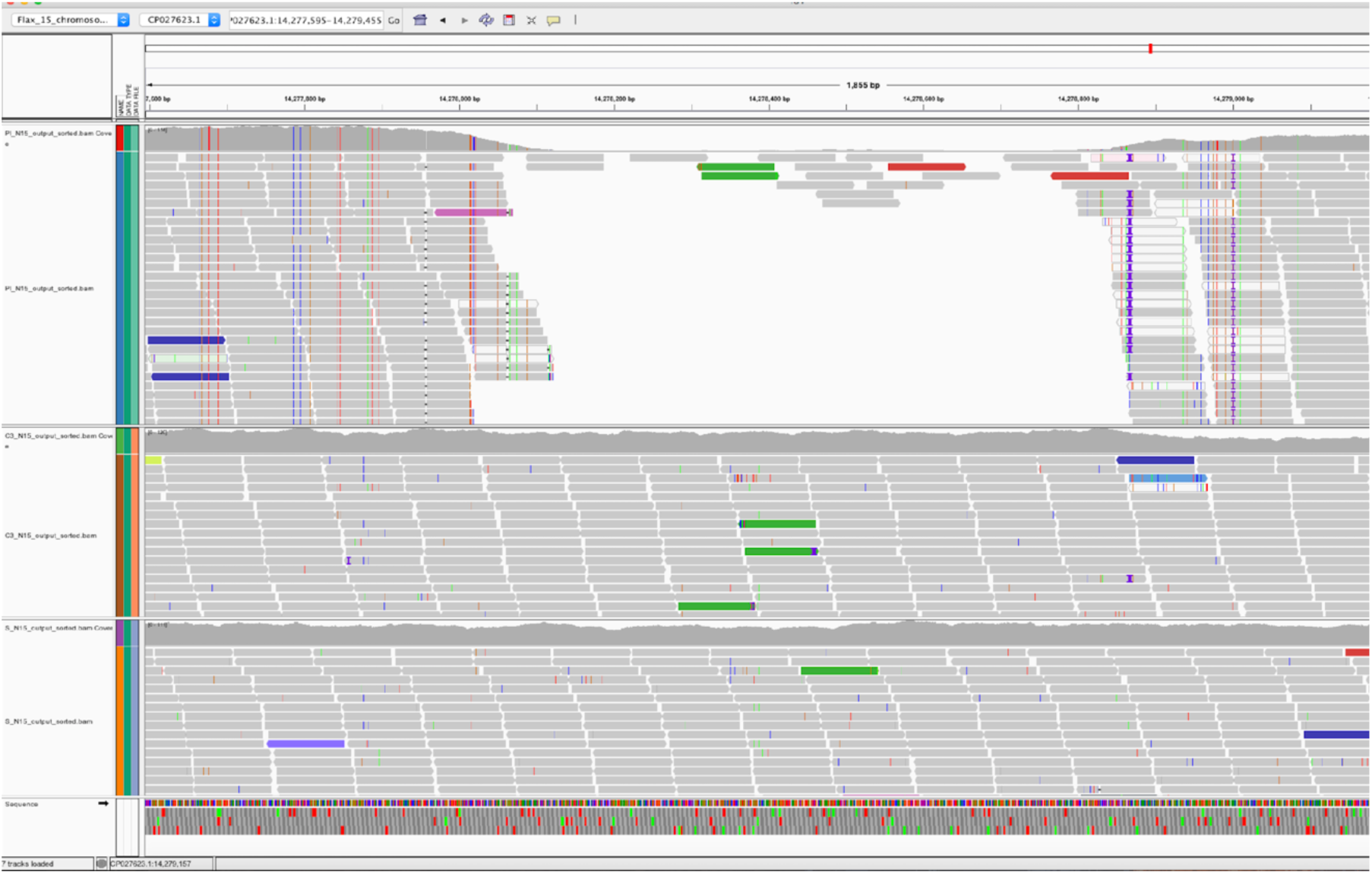
IGV picture at site 6. There is a gap and multiple SNPs sites between Bethune and Pl. While C3 and S are very similar to Bethune.

**Figure 25.**
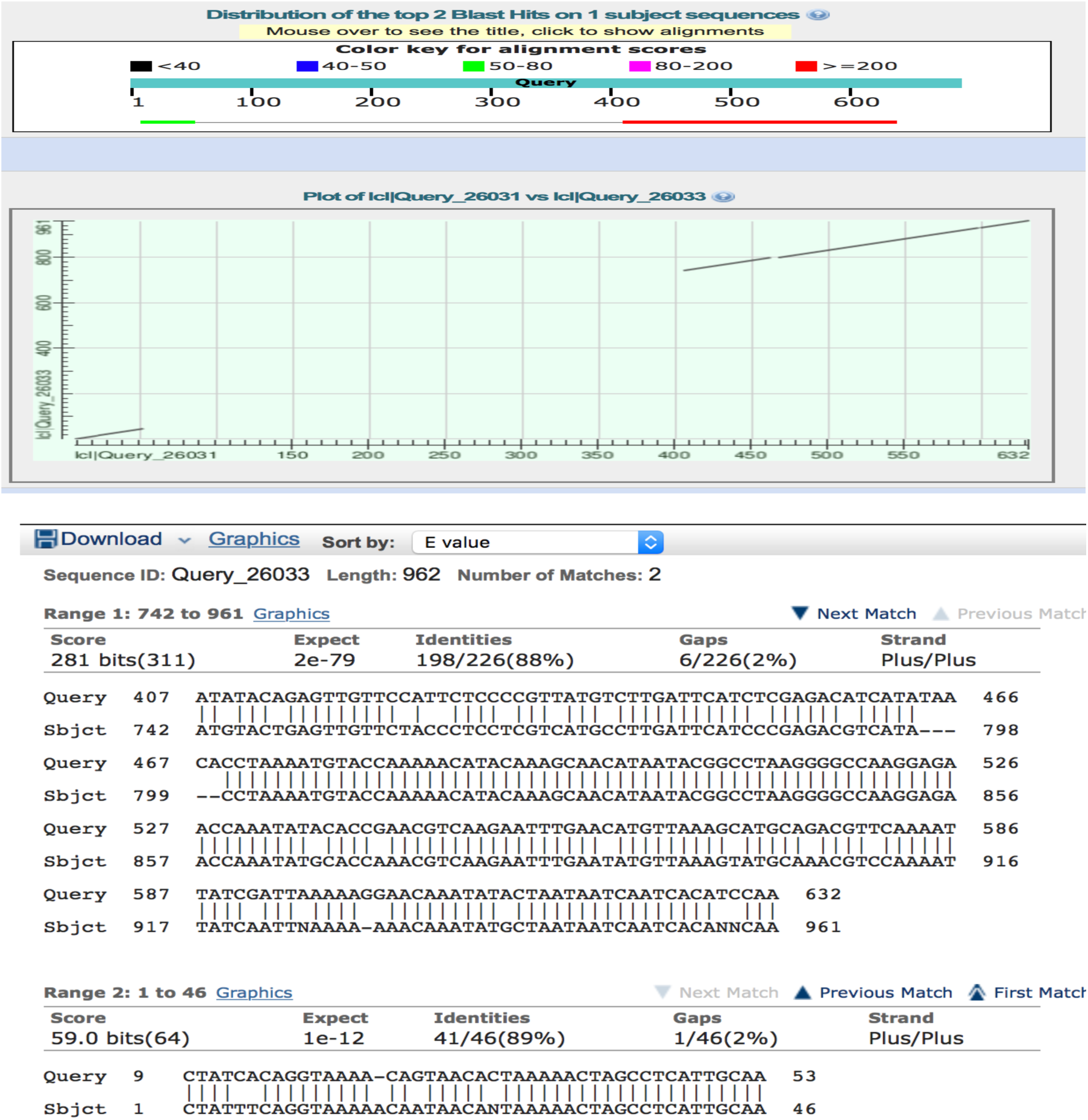
BLAST result of PCR reaction amplified by primer 6 of two samples that show variation on a gel.

**Figure 26.**
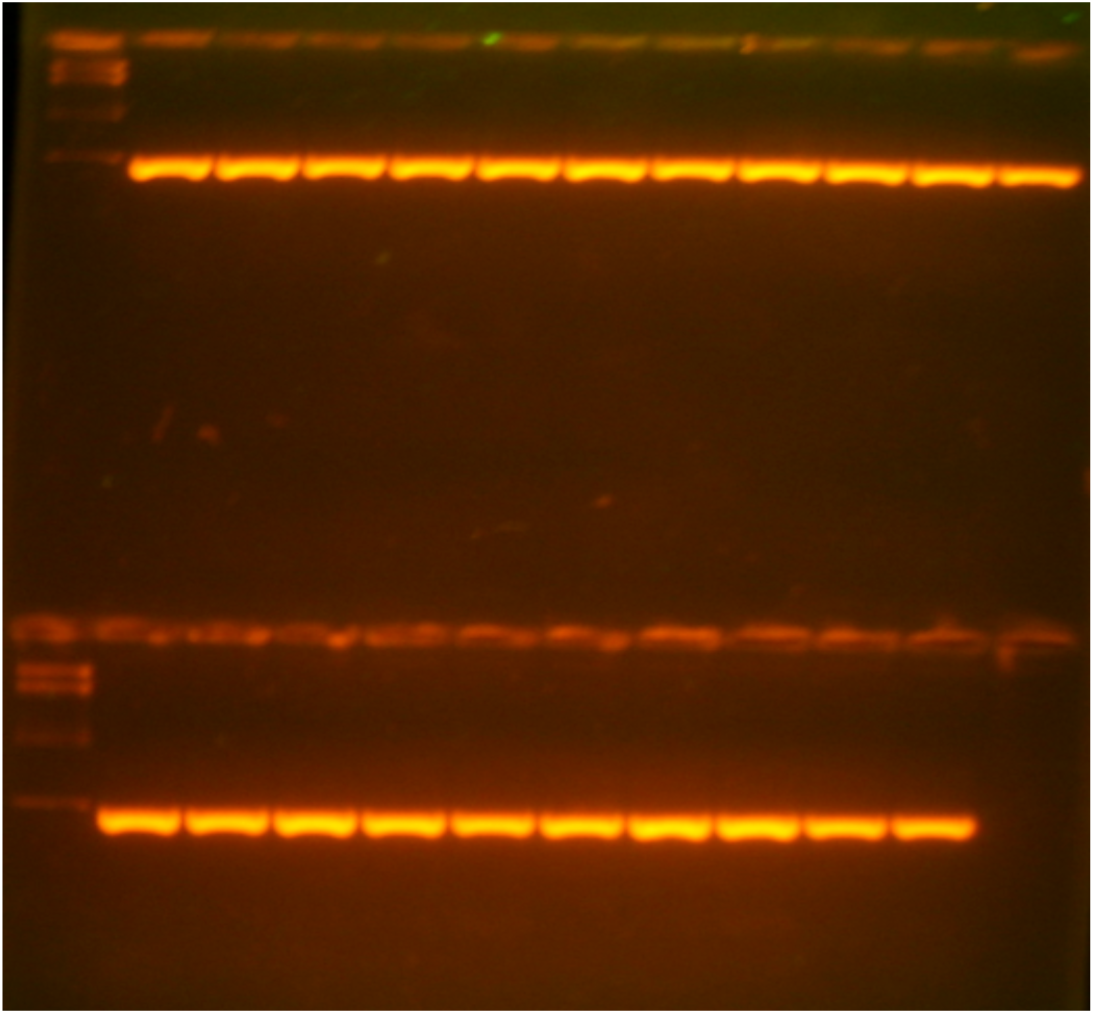
PCR reaction amplified by primer 34. 1:marker II 2:1wl. 3:1nl 4:2wl 5:2wm 6:2nl. 7:2nm 8:3wl 9:2wm 10:3nl 11:3nm 12:4wl 13: marker II 14:4wm 15:4nl 16:3wl 17:3wm 18:3nl 19:3nm 20:4w 21:4wm 22:4nl. 23:4nm. Gel condition: 1.2% 75V 3hours.

### 3 Samples treated with different nutrition condition

BLAST between 1nl and 2wl (samples show variation on the gel) shows two matches with 117 bases and 17 bases. Query sequence refers to sample “1nl” and subject sequence represents “2wl”. The red frame means the repeat sequence. The “CCAAA” at 58-62 bases of subject sequence shows twice in the query sequence from 776 to 780 and from 14-18.

“Blastn” is used to compare aligned two DNA sequences. Query sequence refers to sample “1nl” and subject sequence represents “2wl”. The red frame represents the repeat sequence. The 348-353 bases of query “ACCGAA” sequence show two times in the subject sequence from 345-350 and from 378-383. (Please note that I changed one base from the true DNA in subject sequence to make the second match show, so the second match is not 100%, it is actually 18/19 (95% identities)).

The gap at end of the line shows an overlap suggesting a small insertion happen at this site, which possibly is a transposon because of the overlap.

Query sequence refers to sample “1nl” and subject sequence represents “2wl”. The sequences match each other at start and end with 46 bases and 226 bases. At middle, the DNAs are in different sequence from each other. IGV shows a 800bp gap between them, however both gel and BLAST suggest a 350 real gap between the two genotrophs, which will be further considered in the discussion.

The primer 34 site didn’t show any difference in this set of plants. For the IGV result of primer 34 comparison is shown in Figure 10.

The sample 2wl was collected after 5 weeks growth and shows an extra band of 2/3’ which indicates a preference of LIS-1 insertion in low nutrition condition. For the other samples, one band for 2/pl9 primer and no band for 18a/19 primer which means the LIS-1 insertion is not completely contained. For primer 1,2,6, the sample 2wl and 2wm collected after 5 weeks growth show obvious different than others. They have a smaller band for primer 1 and a larger band for primer 2 and 6. The difference between larger band and smaller band for primer 1 is around 800bp, while 50bp for primer 2 and 300bp for primer 6.

## Discussion

The results bear on the question as to how flax genome can change in particular way to adapt to the new environment and specifically on the process that forms the LIS-1 insertion. This evidence not only contributes to the characterization of large-scale DNA changes, but also provides new data on how some of these particular genomic changes are regulated. In light of these findings possible next steps are suggested.

Depending on previous LIS-1 model (Figure 4) and Cullis (2017) and Chen (2005), when LIS-1 is fully inserted, 2/3’ and 18a/19’ bands show up while 2/pl9 band disappears; when LIS-1 is not inserted, 2/3’ and 18a/19’ don’t show band but 2/pl9 has a band. However, the results reported here are, in some ways similar to those reported in Chen et al. (2005) when amplification across the ends of LIS-1 can occur while the uninserted site can still be amplified. The data for the third set plants show the replacement of the uninserted site with LIS-1. However, for the first and second set of plants, those three pairs of primers did not have the expected relationship (First, second and third sets of plants were described in the Materials and Methods section). The three primers can all amplify or lose a band from the same DNA. In addition, 2/3’ and 18a/19’ were not consistently found together, that is they were present independently of each other. For the case that they were all amplified, the original DNA could have no insertion, but the 2/3’ and 18a/19’ are formed somewhere else, but not necessarily integrated into the genome, but may have been extrachromosomal. In this scenario, the LIS-1 insertion would be assembled external to the chromosomes and inserted when the assembly is complete. For the case where they all have no amplified fragment, middle part of LIS-1 may have been inserted before the ends, indicating a possible multiple DNA crossover process. For 2/3’ and 18a/19’, they may be independently formed on an extrachromosomal site when a responsive plant experiences stress.

For genomic changes on different branches or leaves, there is no significant rule found about in which part of the plant changes earlier, that is all the variants appear to occur independently. Although the data from figure 1 is not that significant to support that the change is more likely to happen when the plant growth in stress with longer time (top and middle show more change rate), but, they do demonstrate that the genome change can happen at any time during development, since they were seen at bottom, middle and top of the stem. Also, the genomic changes can exist on most of the branches of one plant, none of the branches or some of the branches, which would be consistent with different timing of the genomic alteration. These results suggest that there is no individual control or developmental regulation of where the changes should be made, instead, the variation can be formed anywhere in the plant at anytime, but only if the environment can stimulate the plant response. As a result, the plant can change the DNA of most cells, including flowers and seed, and then the next generation could inherit those genetic changes. This type of genome change reveals the directional and intentional genome arrangement because of the reproducibility of the variants. Also, if stress can cause specific genome changes, what will happen if all genome-changing sites have been altered? The accurate and precise arrangement brings a potential limitation of the genome variation of a line like Pl since C3 and S, would be similar to Bethune when living in low nutrition, but all are uninducible in that low nutrition environment. A hypothetical mechanism for development of these variants could involve RNA, like a RNA retro-transposable element, because DNA change has to be come from one or more templates instead of being randomly made. If some sequences are newly formed, they have to be copied from a brand new nucleotide sequence so small non-coding RNA may be responsible for this due to their mimic character by which the new sequence was made and copied to the original DNA. This may also explain the potential miRNA sequences encoded by LIS-1 (Moss and Cullis, 2012). If this is right, maybe RNA is the involved in facilitating the genomic change. Generally, when cell detects the stress, particular proteins can be activated by the cell pathway involved with cell membrane and second messenger and makes nicks or causes DNA instability at its recognition site like a restriction enzyme. Then small RNAs may anneal to end of the nick and work as a template making new sequence similar to DNA sequence from somewhere else in the genome that is the template or functioning like the CRISPR/cas9 system (Jiang et al., 2013).

For primer 1, 2 and 6, the variations happen on the samples number 2 living in the low nutrition (2wl and 2wm). However, the change mechanism for those three sites may be different. Amplifications using primer 1 results in a 700 base-gap in C3 compare to Bethune and Pl from positions 16209200 to 16209900 bases on chromosome 13 in IGV, which is consistent with the gel band that 2wl and 2wm have around 700 smaller size band than other. The area around the gap appears clean and same as reference from IGV, so it is probably a simple deletion. The BLAST shows a 5 base repeat in a sequence and 5 base overlap in 2wl and 2wm, which is consistent with a transposon footprint. The inserted sequence does have matches to any other part of flax genome. Therefore the primer 1 site variation can be caused by the excision of a transposon, although the sequence of the excised region does not match any transposable element in the NCBI database. Primer 2 contains around 30 bases insertion compared to Pl which is also consistent with the gel picture. The 9kb area around primer 2 from 5800kb to 5809kb on chromosome 1 in IGV shows more SNPs and genome arrangement activities which indicates a potential relation between primer 2 and this entire area. The DNA outside the 9kb area remains identical like 5809kb to 5825kb and 5790kb to 5800kb. Thus, the primer 2 insertion can be a part of a huge genome arrangement from 5800kb to 5809kb. This could be resulted from a big assembled DNA replaces the whole area, or many independent activities happen on this site and cause instability of DNA structure and cause more replication error as SNPs. The BLAST result also shows a 6-base repeat in one sequence but overlapped in another, suggesting a 30-base transposable element footprint when considering the primer variation as an independent change. The primer 6 site shows a 700 insertion in IGV from 14276100 to 14276800 bases on chromosome 13. However, both gel picture and sequencing result identifies a 350 base difference. The 350-base difference is correct, but the 700-base gap is due to an unseen 350bp sequence present in standard Pl genome that is replaced in the genotrophs. So, the Pl has 350-base insertion compared to Bethune and those bases are removed when Pl was induced to C3 and S, that is the original 350 bases insertion is replaced by 700bp fragment which is identical to that present at this position in *Bethune*. This is a more complicated arrangement in that there is an existing fragment in Pl from 53 to 407 in query sequence that is different from reference genome and the net insertion to Pl is actually 332 bases some sequence from 46 to 742 of the subject sequences (Figure 29). The real insertion can be larger than 332bp because the original 354bp existed sequence is also replaced or some other more complex process like two insertion events happen at sides of the 354 existed inserted fragment. Otherwise, this area also shows many polymorphisms just like primer 2 showed. So, potentially, the primer 6 variation is also a part of another larger arrangement unit.

If RNA plays a role in genome variation and DNA variation is limited due to RNA control, the phenotype of a species can be restricted in a range. If same RNAs keeps working on the specific part of genome to cause the same or similar genomic change, there could be a maximum extent of genome change. So, the new species formation would need large scale genome change to create new dimension of the RNA regulation net either by extreme environment or hybridization. The newly formed species will belong to its origin, but it would have new and different chromosomal genome pairs. Not only can it have the phenotype or advantage and function or sensitivity from all the original parents, the new genomes meet and create a new RNA regulation pattern to provide new dimension phenotype which increases the evolution potential and direction. The common genes translating into protein will contribute to the basic organs and cells development as most creatures do and need to be conserved.

For the future work, there still a need to look at the whole genome comparisons and find more variation sites and analyze them in detail about what happens, what is the possible mechanism and how these fragments function. Primer 2 and primer 6 need more work to know if they arrange independently or are involved with other larger fragment arrangements. It is still not clear why a particular sequence is inserted or deleted at specific site. Plasmid (or an extrachromosomal DNA), as a vector that can recombine with the chromosomes, can be a potential genome regulation approach to explain the large fragment insertion or deletion, as could some RNAs. Some of the data still need to be further confirmed to ensure the accuracy. There is also a need to look at DNA structure, DNA expression level, RNA and protein changes and other related work to combine biochemistry, cell biology and genetics to fully understand the meaning of plant genome reaction to itself and how these changes results in different phenotype.

